# The Src and Abl family kinases activate the Spleen Tyrosine Kinase to maximize phagocytosis and *Leishmania* infection

**DOI:** 10.1101/2022.11.14.513575

**Authors:** Imran Ullah, Umaru Barrie, Rebecca M. Kernen, Emily T. Mamula, Francis Tho Huu Khuong, Laela M. Booshehri, Emma L. Rhodes, James M. Bradford, Arani Datta, Dawn M. Wetzel

## Abstract

*Leishmania* spp. are obligate intracellular parasites that must be internalized by phagocytic cells to evade immune responses and cause disease. The uptake of both *Leishmania* promastigotes (insect-stage parasites) and amastigotes (proliferative stage parasites in humans and mice) by phagocytes is thought to be mainly host cell-driven, not parasite-driven. Our previous work indicates that host Src and Abl family kinases facilitate *Leishmania* entry into macrophages and pathogenesis in murine cutaneous leishmaniasis. Here we demonstrate that host spleen tyrosine kinase (SYK) is required for efficient uptake of *Leishmania* promastigotes and amastigotes. A Src family kinase-Abl family kinase-SYK signaling cascade induces *Leishmania* amastigote internalization. Finally, lesion size and parasite burden during *Leishmania* infection is significantly decreased in mice lacking SYK in monocytes or by treatment with the SYK inhibitor entospletinib. In summary, SYK is required for maximal *Leishmania* uptake by macrophages and disease in mice. Our results suggest potential for treating leishmaniasis using host cell-directed agents.

**SUMMARY STATEMENT:** Activation of Spleen Tyrosine Kinase by Src and Abl family kinases is required for maximal *Leishmania* uptake by macrophages and disease in a mouse model of cutaneous leishmaniasis.

## INTRODUCTION

Approximately 350 million people worldwide are at risk for the disfiguring cutaneous or fatal visceral forms of leishmaniasis, which is a neglected tropical disease (WHO, 2022). The drugs currently used to treat leishmaniasis have poor efficacy and/or are highly toxic to human hosts; therefore, new antiparasitics are urgently needed (Wetzel and Phillips, 2018). Leishmaniasis results from infection by obligate intracellular parasites in the *Leishmania* genus. During its life cycle, *Leishmania* alternates between promastigotes, found in sand fly vectors, and amastigotes, found in vertebrate hosts such as humans, dogs, and rodents. In order for infection to be established in humans after a sand fly bite, phagocytic cells (*e.g.*, neutrophils, macrophages [Mϕ]) must ingest promastigotes, using a process that is thought to be primarily driven by the host cell (Ueno and Wilson, 2012). The parasites survive in acidic phagolysosomes, where they differentiate into amastigotes and multiply. If amastigotes exit this compartment, they must be taken up again by phagocytes to survive within their vertebrate hosts (Ueno and Wilson, 2012). *Leishmania* parasites that remain outside of phagocytic cells are thought to either die or be killed by the host immune system.

Multiple protein receptors on the surfaces of phagocytic cells have been implicated in the process of *Leishmania* uptake. For example, promastigotes bind the complement receptor CR3; this process is enhanced when parasites are opsonized (coated) with the terminal component of complement, C3bi (Mosser *et al*., 1992; Puentes *et al*., 1988; Russell and Wright, 1988). Amastigotes are opsonized with immunoglobulin G (IgG) and bind the Fc receptor subclass FcγR (Guy and Belosevic, 1993), which is required for a productive *in vivo* infection (Kima *et al*., 2000). After *Leishmania* binds cell surface receptors, actin-rich phagocytic cups form and surround the parasites, allowing them to be internalized (Lodge and Descoteaux, 2008). However, the host cell signaling process that drives *Leishmania* uptake has not been well-described.

Previously, we implicated host Src family kinases (SFK) as well as Abl family kinases in the process of *Leishmania amazonensis* uptake by Mϕs. The SFK, and specifically Hck, Fgr, and Lyn, are non-receptor tyrosine kinases that had been known to participate in IgG-mediated phagocytosis (Fitzer-Attas *et al*., 2000) after pathogens bind the Fc Receptor (FcR) (Wu *et al*., 2001). Using SFK inhibitors such as SU6656 and primary M<s lacking Hck, Fgr, and Lyn, we demonstrated that SFK were necessary for maximal amastigote uptake by Mϕ, as well as for disease in a mouse model of cutaneous leishmaniasis (Wetzel *et al*., 2016). SFK bind and phosphorylate another set of non-receptor tyrosine kinases, the Abl family kinases Abl (Abl1) and Arg (Abl2) (Mader *et al*., 2011; Plattner *et al*., 2004; Tanis *et al*., 2003); (Bradley and Koleske, 2009). We have shown that Abl and Arg facilitate complementary activities during phagocytosis and *Leishmania* uptake by Mϕ (Wetzel *et al*., 2012). Specifically, Mϕ isolated from mice lacking Arg (*Arg^-/-^* Mϕ) exhibited reduced phagocytosis of IgG-coated beads or amastigotes, but had no defects in the uptake of C3bi-coated beads or promastigotes (Wetzel *et al*., 2012). Conversely, Mϕ from mice lacking Abl (*Abl^-/-^* M<) showed defects in the uptake of C3bi-coated beads and promastigotes, but had no defects in the internalization of IgG-coated beads or amastigotes (Wetzel *et al*., 2012). Mϕ lacking both Abl family kinases had no deficiencies in C3bi or IgG-mediated uptake beyond that seen for Mϕs lacking the single relevant kinase (Wetzel *et al*., 2012; Wetzel *et al*., 2016). Mice lacking Abl or Arg, or treated with the Abl family kinase inhibitor imatinib, had smaller lesions and lower parasite burden in a cutaneous mouse model of leishmaniasis (Wetzel *et al*., 2012). Finally, treatment of *Leishmania*-infected mice with bosutinib, which inhibits both Abl family kinases and SFK, led to greater decreases in lesion size and parasite burden than were seen with inhibition of either kinase family alone (Wetzel *et al*., 2016).

Spleen Tyrosine Kinase (SYK) is a cytoplasmic tyrosine kinase with 10 tyrosines that can be phosphorylated to participate in signal transduction from immune receptors such as FcR (Furlong *et al*., 1997). Mϕ derived from mice conditionally lacking SYK in immune cells are defective in several FcR-induced signaling events (Bonnerot *et al*., 1998; Bradshaw, 2010; Lowell, 2011). These Mϕ also have deficits in IgG-mediated phagocytosis; they form actin-rich phagocytic cups but cannot close them around pathogens (Crowley *et al*., 1997). However, these M< respond normally to lipopolysaccharide (LPS), indicating that there are no deficiencies in activation (Crowley *et al*., 1997). Some laboratories have suggested that SYK inhibition does not affect C3bi-mediated phagocytosis (Kiefer *et al*., 1998), but others have shown significant defects in this process (Tohyama and Yamamura, 2006; Walbaum *et al*., 2021). There do not appear to be SYK homologs in *Leishmania* (Baker *et al*., 2021; Parsons *et al*., 2005).

Prior studies suggest that kinases previously implicated in *Leishmania* internalization also may modulate SYK-mediated processes in related biological systems. For example, Mϕ derived from mice deficient for Hck, Fgr, and Lyn exhibited decreased SYK activation when FcR was crosslinked by antibodies (Crowley *et al*., 1997). In addition, loss of Abl/Arg activity limited SYK phosphorylation after FcR crosslinking with antibodies, and Arg kinase phosphorylated SYK in kinase assays with purified proteins (Greuber and Pendergast, 2012). Based on these prior findings, we hypothesized that host cell SYK would be activated by SFK and Abl family kinases during the uptake of *Leishmania* by M<, and that SYK activity would be required for pathogenesis in a mouse model of leishmaniasis.

Here, we demonstrate that host cell SYK facilitates both C3bi- and IgG-mediated phagocytosis and promotes the internalization of *Leishmania* promastigotes and amastigotes. A SFK-Abl family kinase-SYK pathway is required for maximal internalization of *Leishmania* amastigotes by Mϕ. In addition, mice lacking SYK in monocytes, as well as mice treated with the SYK kinase inhibitor entospletinib, have significantly reduced lesion severity and parasite burden in a mouse model of cutaneous *Leishmania* infection. Our results provide proof-of-concept that leishmaniasis could potentially be treated with agents directed at host cell kinases such as SYK.

## RESULTS

### Entospletinib decreases complement and immunoglobulin-mediated phagocytosis

Previously, we have shown that SFK inhibitors and Abl family kinase inhibitors decrease the uptake of *Leishmania* parasites (Wetzel *et al*., 2012; Wetzel *et al*., 2016). Others have suggested that signaling during phagocytosis by these two kinase families can be relayed to SYK (Crowley *et al*., 1997; Greuber and Pendergast, 2012). Therefore, we chose to explore the role of SYK in phagocytic model systems and *Leishmania* uptake by M<.

Entospletinib is a SYK inhibitor that has an IC_50_ of 7.7 nM in purified kinase assays (Ramanathan *et al*., 2017). Given its recent favorable results in a human phase 1b/2 trial for acute myeloid leukemia (Walker *et al*., 2020), it is an attractive tool for further study. However, its effects on phagocytosis had not been documented previously. For this reason, we first determined whether entospletinib treatment of phagocytic cells affected immunoglobulin-mediated phagocytosis. We treated RAW 264.7 cells, a murine Mϕ-like cell line, with increasing concentrations of entospletinib or DMSO for 2 hours (h) and incubated them for 30 minutes (m) with beads that had been coated with IgG. We found that entospletinib, in a concentration-dependent manner, decreased the uptake of IgG-coated beads, as demonstrated by the phagocytic index or number of internalized beads per 100 Mϕs (PI, Fig. S1A). Based on this titration, we selected a concentration of 1 μM for further studies.

Whether SYK plays a role in C3bi-mediated phagocytosis has been a subject of debate (Kiefer *et al*., 1998; Tohyama and Yamamura, 2006; Walbaum *et al*., 2021). To determine whether entospletinib treatment of host cells affected C3bi and IgG-mediated phagocytosis, we treated RAW 264.7 cells with 1 μM entospletinib or DMSO for 2 h and incubated them for 30 m with beads that had been coated with C3bi or IgG. We found that 1 μM entospletinib decreased both C3bi-coated and IgG-coated bead uptake by RAW 264.7 cells by 44 ± 6% and 67 ± 8% (mean ± standard error [SE]), respectively (Fig. 1A, B). Since RAW 264.7 cells are a cancer cell line that has been virally transformed, these results were confirmed with primary cells: bone marrow-derived Mϕs (BMDM) isolated from mouse tibias/femurs (Fig. 1C). The decreases in C3bi- and IgG-mediated phagocytosis due to entospletinib treatment specifically resulted from defects in internalization, as entospletinib did not affect the total (internal + external) number of C3bi- and IgG-coated beads normalized to the number of RAW 264.7 cells (Fig. 1D) or BMDM (Fig. 1E).

**Fig. 1.**
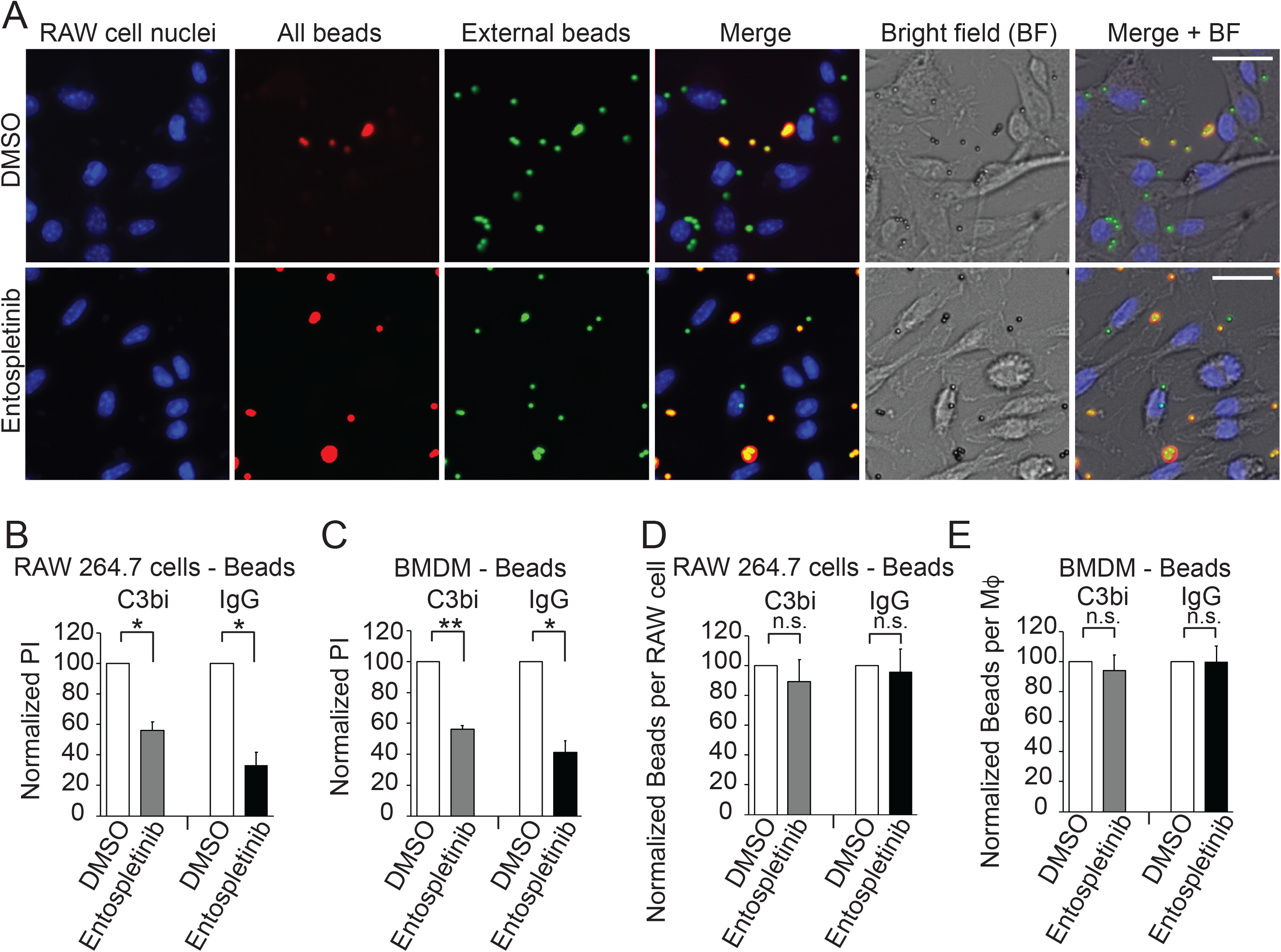
The SYK inhibitor entospletinib decreases C3bi and IgG-mediated phagocytosis. M<s were treated with 1 μM entospletinib or DMSO for 2 h and incubated with C3bi or IgG-coated beads for 30 m. A multi-colored immunofluorescence assay was used to distinguish between extracellular (red + green) and intracellular (green) beads (see methods for details). Nuclei are labeled with Hoescht 33258. (A) Image of anti-P8 C3bi-opsonized bead uptake by RAW 264.7 cells treated with DMSO (top) or entospletinib (bottom). Scale bar = 10 μm. In the top panel, there are portions of 12 M< nuclei visible, with 21 total beads and 7 external beads (*i.e.,* 14 internalized beads), suggesting a phagocytic index (PI) of 117. In the bottom panel, 11 M< nuclei are seen, with 20 total beads and 16 extracellular beads (*i.e.,* 4 intracellular amastigotes), suggesting a PI of 36. (B) Quantification of the effects of entospletinib on C3bi-coated and IgG-coated bead uptake by RAW 264.7 cells. Shown is the mean PI ± SE for RAW cells incubated in 1 μM entospletinib and allowed to take up C3bi-coated (left) or IgG-coated beads (right), normalized to RAW cells incubated in an equivalent volume of DMSO and the same number of beads (shown as PI = 100%). (C) Enumeration of the effect of entospletinib on C3bi-coated and IgG-coated bead uptake by BMDM. Shown is the mean PI ± SE, calculated through the methodology described in (B). (D, E) Entospletinib does not affect the total number of C3bi- and IgG-coated beads bound to (D) RAW 264.7 cells and (E) BMDM. Shown is the total number of beads per M< ± SE after M< were incubated in 1 μM entospletinib and allowed to take up C3bi-coated (left) or IgG-coated beads (right), normalized to M< incubated in an equivalent volume of DMSO and beads. For B-E: n = 3 separate experiments; \**p* < 0.05, \*\**p* < 0.01 (one sample t-test).

### SYK inhibition decreases *Leishmania* promastigote and amastigote uptake by M<s

We can grow both promastigotes and amastigotes of *L. amazonensis* without mammalian host cells (axenic cultures), using a supplemented growth media at pH 7.2 for promastigotes or pH 5 for amastigotes (Hodgkinson *et al*., 1996). To test whether entospletinib treatment also affected the internalization of promastigotes and axenic amastigotes, we again pre-treated Mϕs with 1 μM entospletinib or DMSO, but now incubated them with C3bi-coated *L. amazonensis* promastigotes or IgG-coated axenic amastigotes. A representative image of amastigote uptake ± entospletinib is shown in Fig. 2A. We found that 1 μM entospletinib decreased promastigote and amastigote uptake by RAW 264.7 cells (by 39 ± 3% and 54 ± 5%, respectively; Fig. 2B) and BMDM (by 49 ± 6% and 66 ± 6%, respectively; Fig. 2C). Titration of entospletinib demonstrated that promastigote uptake and amastigote uptake by RAW 264.7 cells was decreased in a concentration-dependent manner (Fig. S1B and C, respectively). Entospletinib did not affect the total number of promastigotes or amastigotes that bound RAW 264.7 cells (Fig. 2D) or BMDM (Fig. 2E).

**Fig. 2.**
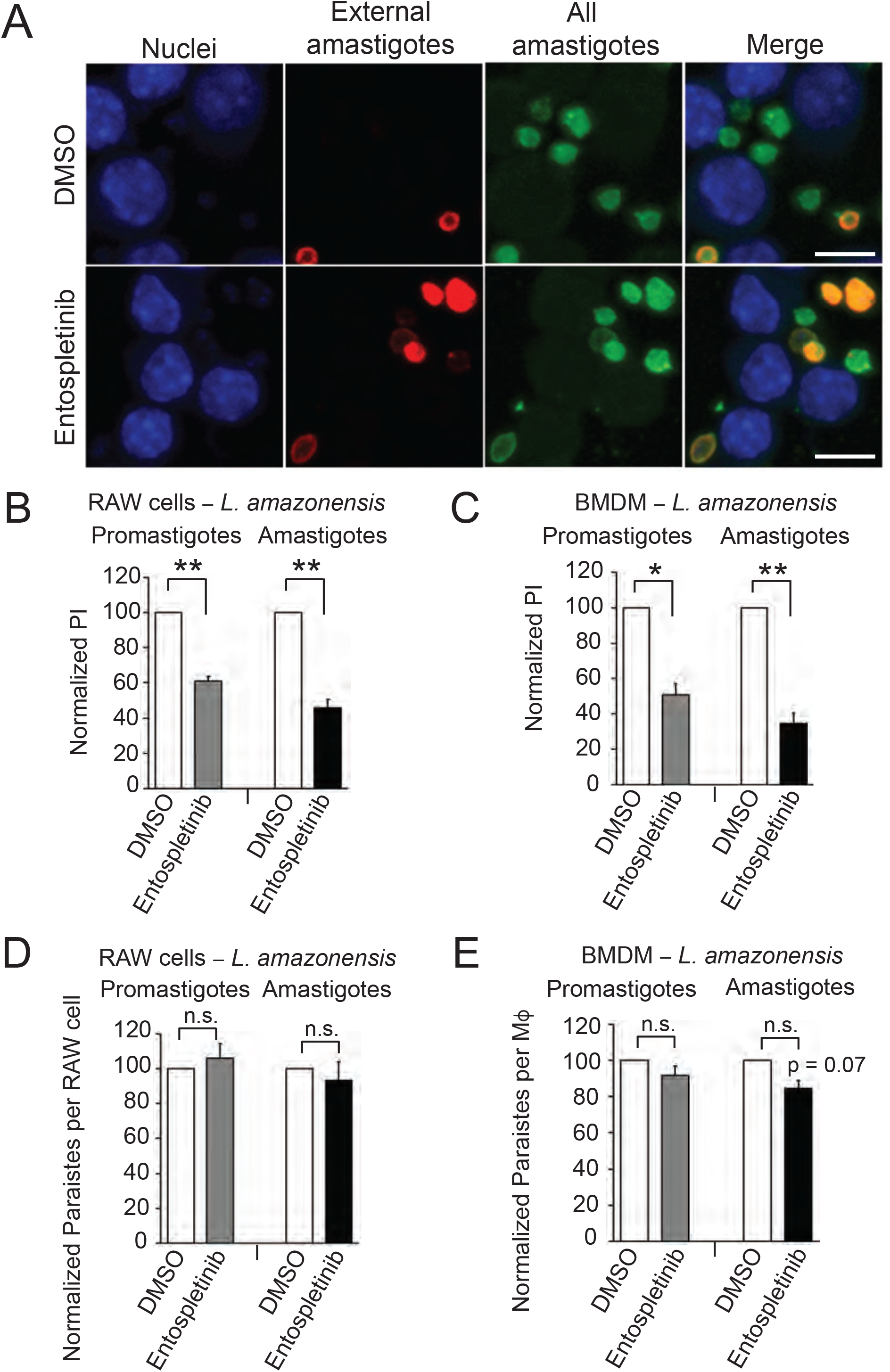
The SYK inhibitor entospletinib decreases *Leishmania* uptake. M< were treated with entospletinib or DMSO for 2 h and incubated with C3bi-coated *L. amazonensis* promastigotes or IgG-coated *L. amazonensis* amastigotes for 30 m. Immunofluorescence was used to distinguish between extracellular (red + green) and intracellular (green) promastigotes or amastigotes. M< nuclei are labeled with Hoescht 33258. (A) Image of anti-P8 IgG-opsonized amastigote uptake by RAW 264.7 cells treated with DMSO (top) or entospletinib (bottom). Scale bar = 5 μm. In the top panel there are portions of 5 M< nuclei visible, with 7 total amastigotes and 2 external amastigotes (*i.e.,* 5 internalized amastigotes), suggesting a PI of 100. In the bottom panel, 4 M< nuclei are seen, with 7 total amastigotes and 5 extracellular amastigotes (*i.e.,* 2 intracellular amastigotes), suggesting a PI of 50. (B, C) Quantification of the effects of entospletinib on promastigote or amastigote uptake by (B) RAW 264.7 cells and (C) BMDM. Shown is the PI ± SE for M< incubated in 1 μM entospletinib and allowed to take up promastigotes (left) or amastigotes (right), normalized to M< incubated in an equivalent volume of DMSO and the same number of beads. (D, E) Entospletinib does not affect the total number of promastigotes and amastigotes bound to (D) RAW 264.7 cells and (E) BMDM. Shown is the total number of parasites per M< ± SE after M< were incubated in 1 μM entospletinib and allowed to take up promastigotes (left) or amastigotes (right), normalized to M< incubated in an equivalent volume of DMSO and the same number of parasites. n = 3 separate experiments; \**p* < 0.05, \*\**p* < 0.01 (one sample t-test).

To ensure that entospletinib did not have deleterious effects on *L. amazonensis* amastigotes, we incubated axenic amastigotes with entospletinib for 72 h and assessed parasite survival with alamarBlue® (Ullah *et al*., 2020). Entospletinib’s EC_50_ against axenic amastigotes in this assay was > 20 μM. The CC_50_ for RAW 264.7 cells over 72 h was 3.3 ± 1.2 μM (3 biological replicates performed; see representative log concentration response curves in Fig. S1D). These studies indicated that entospletinib did not kill either *Leishmania* parasites or host cells over the time period and concentrations studied in these uptake assays. To test whether entospletinib affected the intracellular survival of *L. amazonensis* amastigotes post-internalization, we engineered *L. amazonensis* parasites that expressed mNeonGreen (see methods) and allowed these amastigotes to be taken up by RAW 264.7 cells. 24 h later, we added DMSO or 0.5 μM entospletinib (a concentration where effects on host cells were minimal) for 72 h. We found no apparent defects in the ability of amastigotes to survive within 0.5 μM entospletinib-treated RAW 264.7 cells compared to DMSO-treated RAW 264.7 cells over 72 h (Fig. S1E).

We next explored the mechanism by which entospletinib affected the internalization of *Leishmania* parasites. First, we determined that extending uptake over longer periods (up to 3 h) did not enable treated RAW 264.7 cells to overcome entospletinib-mediated defects in *Leishmania* internalization (promastigotes shown in Fig. S1F; amastigotes shown in Fig. S1G). To more specifically characterize the effects of entospletinib on phagocytic cup formation, we performed an uptake assay with amastigotes as described above, then incubated samples with far-red phalloidin. As has previously been described for SYK inhibition, we found using confocal microscopy that phagocytic cups seemed unable to close in entospletinib-treated RAW 264.7 cells (Fig. S2), particularly since there were no differences in the numbers of cups seen between DMSO and entospletinib-treated M<s (Fig. S3A). In addition, actin staining in phagocytic cups was brighter on-average in entospletinib-treated compared to DMSO-treated RAW 264.7 cells (Fig. S3B), suggesting a direct effect on actin polymerization during phagocytic cup formation.

### Host cell SYK is required for efficient phagocytosis and *Leishmania* uptake by M<s

Given that kinase inhibitors can potentially have off-target effects, and could also be acting upon either parasites or host cells, we next specifically tested the role of host cell SYK during *Leishmania* uptake by Mϕs. Using BMDMs isolated from mice lacking SYK in the monocyte lineage (*Syk^flox/flox^* LysM Cre+, abbreviated as *Syk^-/-^* when discussing isolated BMDM), we found that *Syk^-/-^* BMDM demonstrated decreases in promastigote (Fig. 3A, B) and amastigote uptake (Fig. 3B), by 38 ± 5% and 73 ± 3%, respectively. The total number of promastigotes or amastigotes bound to BMDM was not affected (Fig. 3C). *Syk^-/-^* BMDM also exhibited defects in C3bi and IgG-mediated phagocytosis of coated beads (which were decreased by 43 ± 3% and 65 ± 10%, respectively; Fig. 3D, E).

**Fig. 3.**
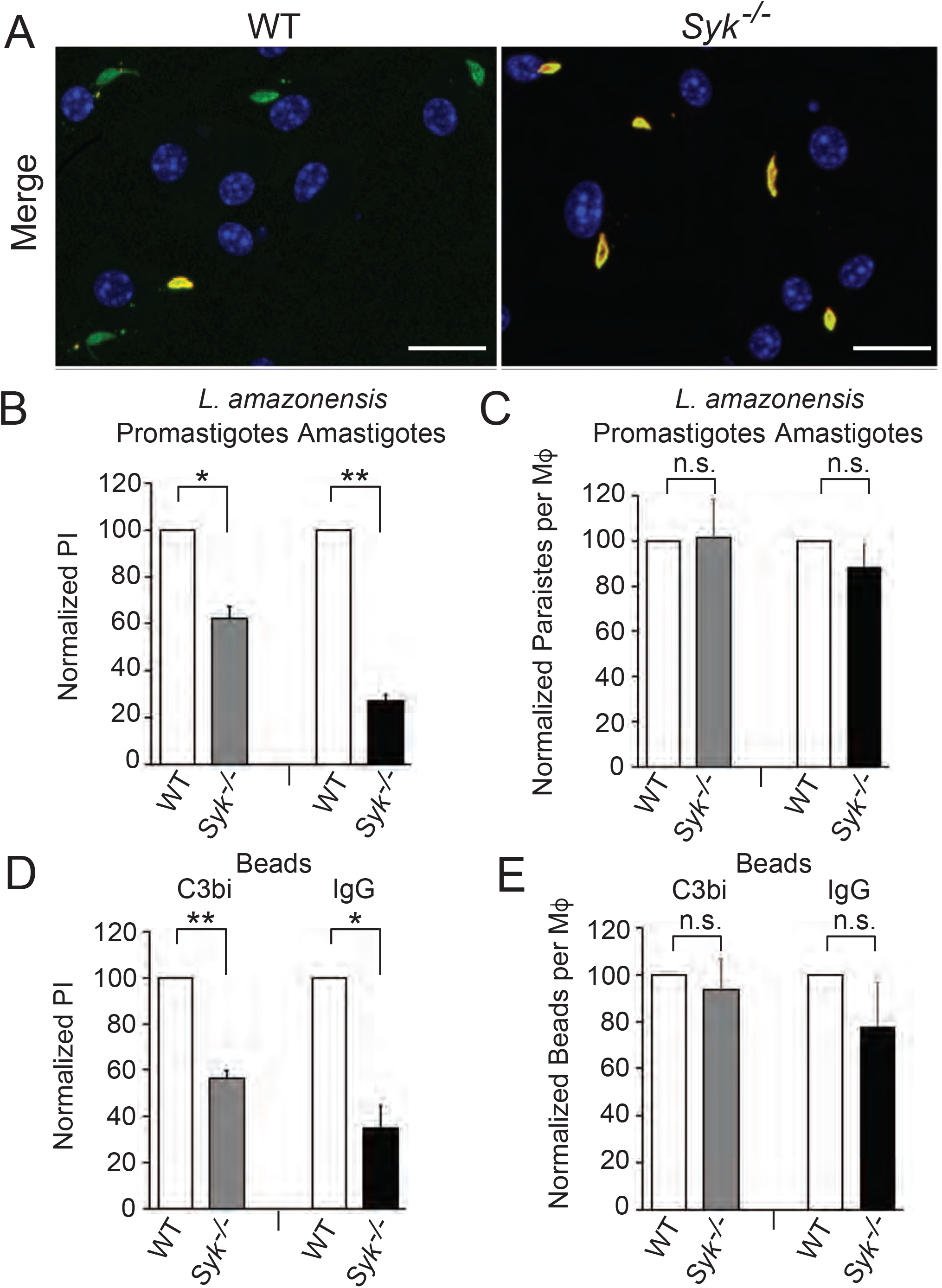
BMDM lacking SYK exhibit defects in C3bi- and IgG-mediated phagocytosis and *L. amazonensis* uptake. (A) Merged image of promastigote uptake by WT BMDM (left) or *Syk^flox/flox^* LysM Cre+ BMDM (*Syk^-/-^* BMDM, right), using the methodology described in Fig 2. Scale bar = 10 μm. On the left of this representative image, there are 8 full M< nuclei visible, with 5 total promastigotes and 4 internalized promastigotes, suggesting a PI of 50. On the right, 7 M< nuclei are seen, with 5 total promastigotes and 0 intracellular promastigotes. (B) *Syk^-/-^* BMDM exhibit defects in promastigote and amastigote uptake. BMDM were incubated with *L. amazonensis* promastigotes or amastigotes as in Fig 2. Graph shows the mean PI ± SE for promastigotes and amastigotes for *Syk^-/-^* BMDM, normalized to WT BMDM, using the methodology described in Fig. 2. (C) Mean normalized total number of promastigotes or amastigotes bound to *Syk^-/-^* BMDM, normalized to WT BMDM, ± SE. (D) *Syk^-/-^* BMDM are less able to undergo C3bi and IgG-mediated phagocytosis. BMDM were incubated with C3bi or IgG-coated beads as in Fig. 1. Shown is the mean PI ± SE for beads taken up by *Syk^-/-^* BMDM, normalized to WT BMDM (using the methodology described in Fig. 1). (E) Mean normalized total number of C3bi or IgG-coated beads bound to *Syk^-/-^* BMDM, normalized to WT BMDM, ± SE. n = 3 separate experiments; \**p*<0.05, \*\**p*<0.01 (one sample t-test).

### *Leishmania* stimulates SFK-Arg-SYK signaling in Mϕs to stimulate uptake

We next explored whether *L. amazonensis* employs signaling by the adhesion molecule PSGL1 (CD162) to facilitate uptake. PSGL-1 has not been shown to be a receptor for *Leishmania* uptake, but it is known to activate SYK signaling via interactions with FcR (Zarbock *et al*., 2008). We studied both of these possibilities using an antibody known to block PSGL-1 signaling (Zanardo *et al*., 2004), which should also prevent *Leishmania* from directly interacting with PSGL-1. We found that this antibody did not affect *Leishmania* internalization (Fig. S3C). In addition, entospletinib still prevented *Leishmania* uptake when added alongside the antibody (Fig. S3D).

SFK and Abl family kinases have been shown to partly signal through SYK in previous studies (Greuber and Pendergast, 2012). We have shown that Hck/Fgr/Lyn specifically activates Abl2/Arg to facilitate amastigote uptake by Mϕs (Wetzel *et al*., 2012; Wetzel *et al*., 2016). SFK are not required for C3bi-mediated phagocytosis in our system (Wetzel *et al*., 2016). Abl1, but not SFK or Arg, is required for efficient promastigote uptake (Wetzel *et al*., 2012; Wetzel *et al*., 2016). Therefore, we hypothesized that an FcR-SFK-Arg-SYK signaling pathway would be required for amastigote uptake, and a CR3-Abl-SYK signaling pathway would be responsible for promastigote uptake. To test this hypothesis, we performed experiments using combinations of kinase inhibitors and activators (Mohamed *et al*., 2022). Treating *Syk^-/-^* BMDM with imatinib to inhibit Abl family kinases did not further decrease *Leishmania* promastigote uptake, and similarly, treating *Syk^-/-^* BMDM with SU6656 to inhibit SFK or imatinib to inhibit Abl family kinases did not further diminish *Leishmania* amastigote uptake (Fig. 4A). Adding entospletinib neither further decreased promastigote uptake by BMDM lacking Abl (*Abl^flox/flox^* LysM Cre+ BMDM; abbreviated in figure as *Abl^-/-^*) nor further impaired amastigote uptake by BMDM lacking Hck, Fgr, and Lyn (*HFL^-/-^* BMDM) or Arg (*Arg^-/-^*BMDM) kinases (Fig. 4B). In addition, the Arg/Abl activator DPH did not rescue the uptake of *Leishmania* promastigotes or amastigotes by *Syk^-/-^* BMDM (Fig. 4C).

**Fig. 4.**
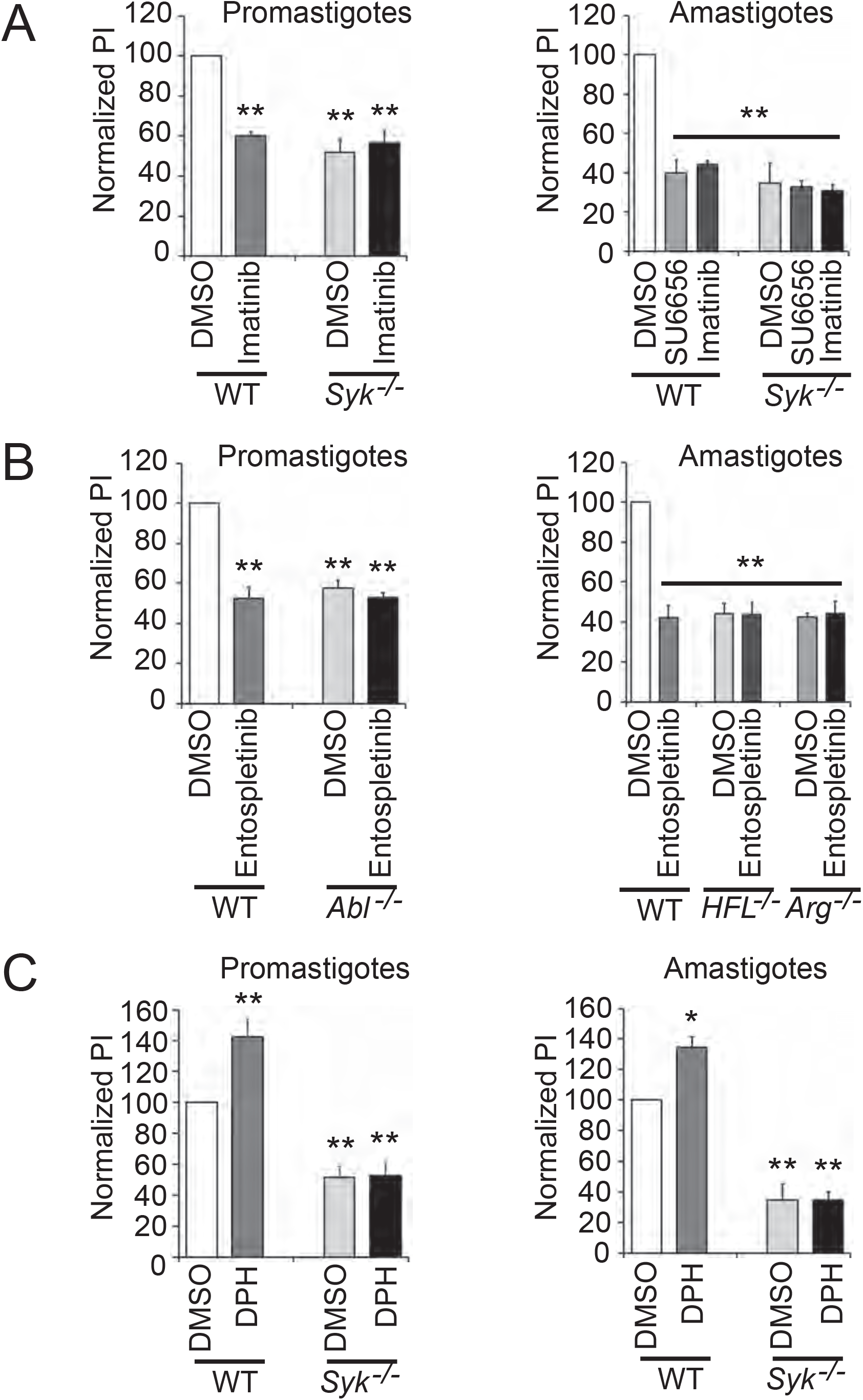
A SFK-Abl/Arg-SYK signaling pathway facilitates *Leishmania* uptake. (A) Treating *Syk^-/-^* BMDM with SU6656 or imatinib does not further diminish *Leishmania* promastigote (left) or amastigote uptake (right). (B) Entospletinib has no effect on promastigote uptake by BMDM lacking Abl (left) or amastigote uptake by BMDM lacking Hck, Fgr, and Lyn or Arg kinases (right). (C) The Arg/Abl activator DPH does not rescue *Leishmania* promastigote (left) or amastigote uptake (right) in *Syk^-/-^* BMDM. For all categories, BMDM of the genotypes shown were incubated with DMSO, SU6656, imatinib, entospletinib, or DPH as indicated and allowed to take up promastigotes or amastigotes. PIs were calculated and values were normalized to WT, DMSO treated BMDM (100%). Mean ± SE shown. n = 3 separate biological experiments; \**p*<0.05, \*\**p*<0.01 by ANOVA compared to WT DMSO treated M<.

We then investigated whether SFK and Abl family kinases were essential for SYK activation during amastigote uptake. We chose to focus on amastigotes as the more biologically relevant life cycle stage in human infection. We allowed M<s to begin internalizing amastigotes and performed immunoblots for phosphorylated SYK (pSYK), normalized to total SYK levels. We found that pSYK levels were decreased after inhibition of SFK (using SU6656) or Abl family kinases (using imatinib), while activating Abl family kinases using DPH increased Syk phosphorylation (Fig. 5A, B; see Fig. S4 for full Western blot images and additional examples). Previously, we have shown that SFK and Abl family kinases are required for effective phosphorylation of their substrate CrkII (*i.e.*, pCrk) during amastigote uptake by Mϕs (Wetzel *et al*., 2012; Wetzel *et al*., 2016). To determine whether SYK was required for this phosphorylation event, we probed for pCrk induction upon amastigote uptake by entospletinib-treated Mϕs. We found lower levels of pCrk in entospletinib-treated Mϕs undergoing amastigote uptake when compared to DMSO controls (Fig. 5C, D; Fig. S4).

**Fig. 5.**
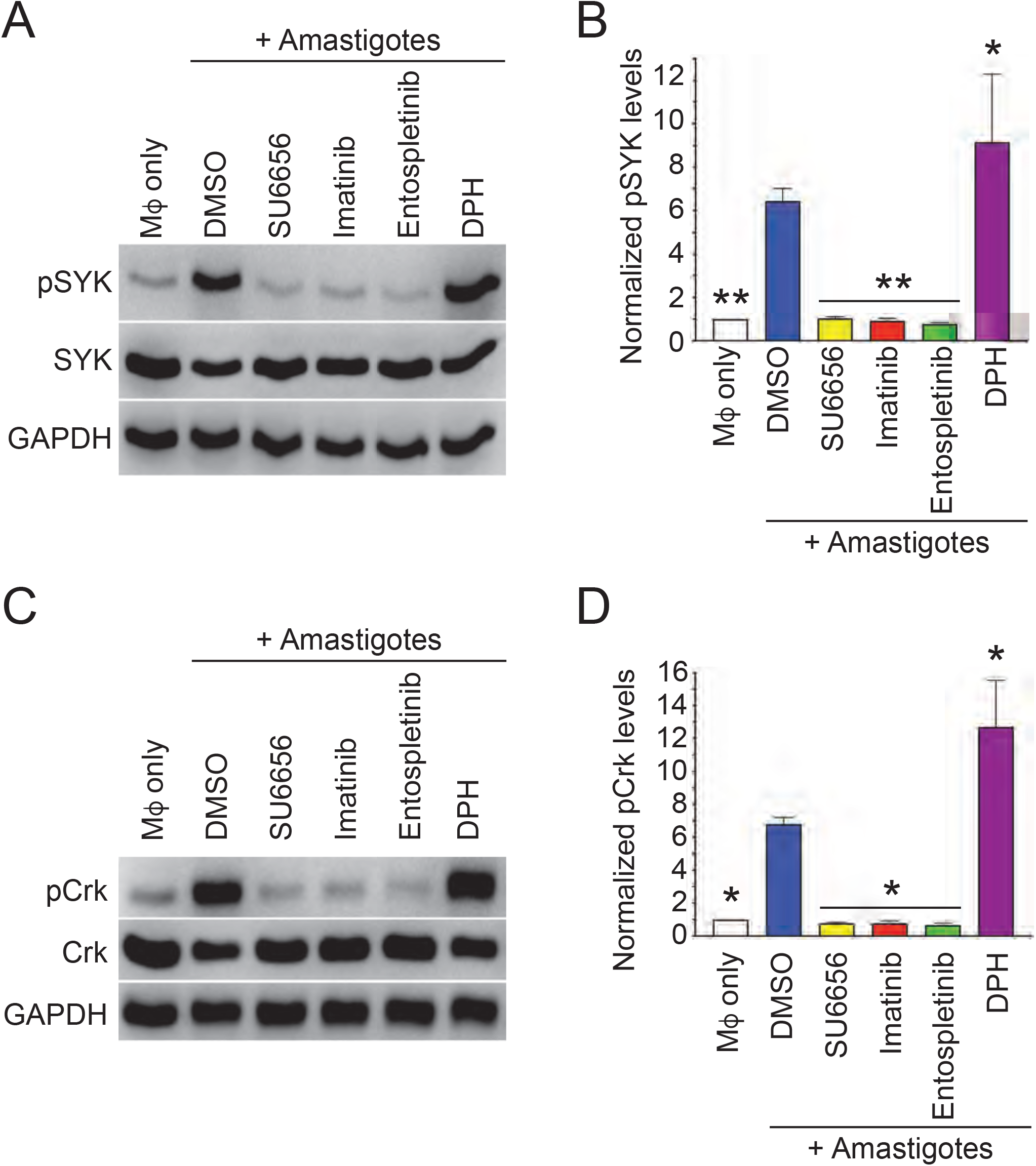
SYK is activated by SFK/Arg and phosphorylates downstream effectors during amastigote uptake. (A, B) SYK phosphorylation is decreased by inhibition of SFK or Abl family kinases and increased by activating Abl family kinases. RAW 264.7 cells treated with DMSO or inhibitors were incubated with IgG-coated amastigotes (*i.e.*, + amastigotes) for 15 m before processing for immunoblotting. (A) Representative immunoblot of pSYK (top) and total SYK (middle) in DMSO-treated RAW 264.7 cells (± amastigotes) and amastigote-stimulated SU6656, imatinib, entospletinib, or DPH-treated RAW 264.7 cells. GAPDH (bottom) is used as an additional loading control. (B) Graph presents relative pSYK levels, normalized to SYK levels, among RAW 264.7 cell categories shown in (A). pSYK/SYK levels for M<s only (*i.e.*, no amastigotes) were normalized to 1. (C, D) Phosphorylation of the SFK/Abl/Arg substrate CrkII (pCrk) induced upon amastigote uptake is decreased in entospletinib-treated M<s. (C) Representative immunoblot of pCrk (top) and total CrkII (middle) in DMSO-treated RAW 264.7 cells (± amastigotes) and amastigote-stimulated SU6656, imatinib, entospletinib, or DPH-treated RAW 264.7 cells. GAPDH is used as an additional loading control. (D) Relative levels for pCrk, normalized to CrkII, graphed as in B. For all categories, n = 4 separate biological experiments; mean ± standard deviations shown. \**p* < 0.05, \*\**p* < 0.01 by ANOVA compared to pSYK or pCrk levels in WT DMSO-treated M< incubated with amastigotes.

### SYK inhibition reduces lesion size and parasite burden in cutaneous leishmaniasis

Knowing that SYK was required for efficient uptake of *Leishmania* by M<s, we investigated whether its activity was required for the manifestations of cutaneous leishmaniasis in mice. We infected *Syk^flox/flox^* LysM Cre+ mice and WT mice in the right hind foot with *L. amazonensis* promastigotes and monitored lesion size over time. We found that *Syk^flox/flox^* LysM Cre+ mice developed smaller lesions than WT mice (Fig. 6A), suggesting that SYK activity is necessary for maximal pathogenesis of cutaneous leishmaniasis in the mouse model.

**Fig. 6.**
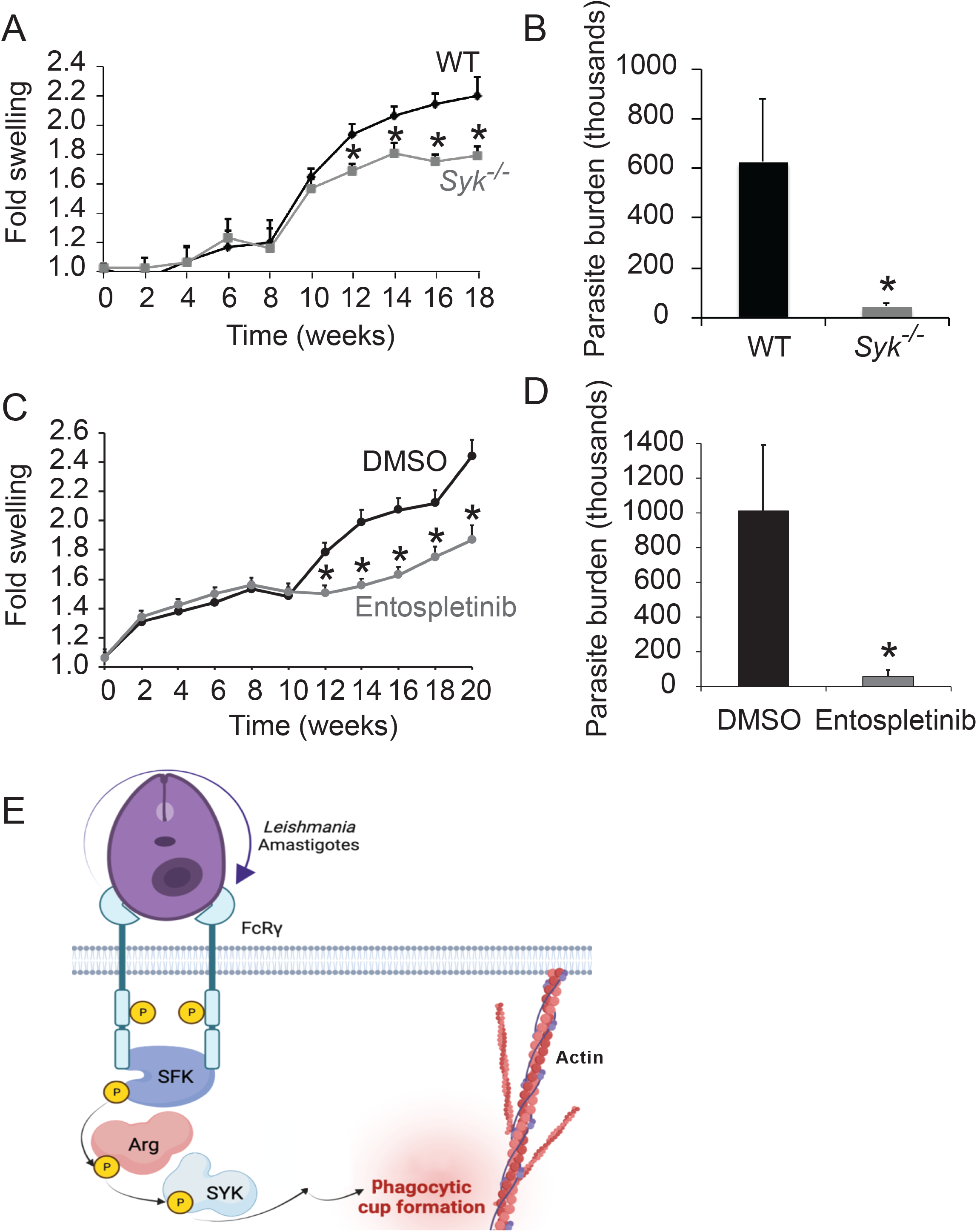
SYK facilitates efficient infection in a mouse model of cutaneous leishmaniasis. (A) Comparison of lesion size in WT versus *Syk^flox/flox^* LysM Cre+ mice. 10 mice per category were infected in the right hind foot with 1 X 10^6^ *L. amazonensis* metacyclic promastigotes and monitored over time. Three experiments were performed; shown is a representative experiment. Graphed are the mean ± SE increase in infected foot width when compared to the uninfected foot. \**p* < 0.05 by ANOVA compared to WT mice. (B) *Syk^flox/flox^* LysM Cre+ mice have a lower parasite burden than WT mice. Plotted is the mean number of parasites isolated from lesions in millions at the termination of (A) at wk 18, ± SE. \**p* < 0.05 by *t*-test. (C) Comparison of lesion size in DMSO/diluent vs. entospletinib-treated mice. Experiment performed as in (A), except that mice were treated with 100 mg/kg/d of entospletinib in their drinking water or the equivalent volume of diluent, starting 4 days prior to infection and continuing throughout the experiment. \**p* < 0.05 by ANOVA compared to DMSO-treated mice. (D) Entospletinib-treated mice have a lower parasite burden than DMSO-treated mice. Shown is the mean number of *L. amazonensis* parasites in millions isolated from infected footpads at the termination of (C) at wk 20, calculated as in (B), ± SE. \**p* < 0.05 by *t*-test. (E) Relationship between FcR signaling, SFKs, Arg, and SYK during *Leishmania* amastigote uptake. Upon FcRγ ligation by amastigotes, host cell Hck, Fgr, and Lyn are activated. These SFKs phosphorylate and activate Arg kinase, which phosphorylates and activates SYK. SYK activates other kinases within the host cell (*e.g.,* CrkII), leading to actin polymerization and parasite uptake. Diagram created with Biorender (https://www.biorender.com).

The smaller lesions seen in *Syk^flox/flox^* LysM Cre+ mice potentially could be the result of differences in their immunological responses to infection. As a simplified paradigm, Th1 responses to leishmaniasis are typically protective, and Th2 responses are generally deleterious to the infected host (Jones *et al*., 1998). Therefore, we isolated draining lymph nodes from infected WT and *Syk^flox/flox^* LysM Cre+ mice and assayed cytokine secretion after stimulation with parasite lysates compared to Concanavalin A (Con A) stimulation (a positive control) or no stimulation (a negative control). Overall, we found limited differences between these two categories of mice (Table 1; Fig. S5A-E), with the exception of IL1β induction, which SYK is known to facilitate (Callaway *et al*., 2015). However, decreases in IL1β have been associated with worsening lesions (Lima-Junior *et al*., 2013). We also did not see any changes in the overall ratio of Th1 vs Th2 cytokine release (Table S1).

**Table 1.**
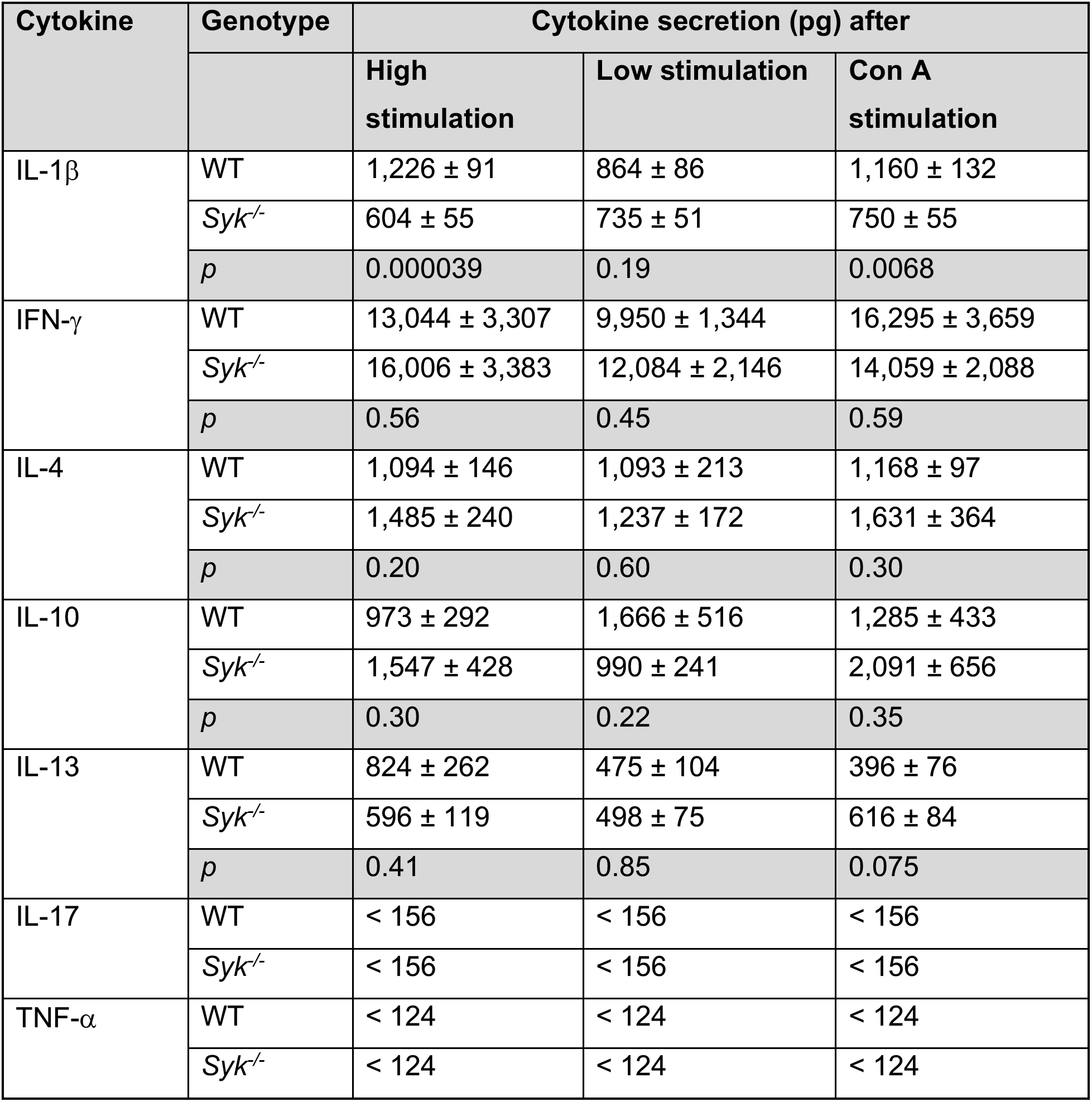
Cytokine secretion in WT vs *Syk^flox/flox^* LysM Cre+ mice. (abbreviated in the table as *Syk^-/-^*). In general, cytokine secretion after *Leishmania* antigen stimulation is not significantly affected in *Syk^flox/flox^* LysM Cre+ mice. The exception is secretion of IL1-β, for which SYK activity is known to be required. Shown are mean ± SE of cytokine profiles of draining lymph nodes isolated from the WT versus *Syk^flox/flox^* LysM Cre+ mice at the end of the experiment shown in Fig 6, with *p* values calculated by Student’s *t*-test. 5 x 10^6^ cells were cultured for 72 h in medium alone (unstimulated, resulting in values that were negligible), or with the addition of lysates made from 2.5 x 10^6^ (high stimulation) or from 5 x 10^5^ (low stimulation) *L. amazonensis* parasites. Concanamycin A (Con A) was used as a positive control for stimulation. Media was harvested and used in ELISAs for the above-listed cytokines as described previously (Wetzel *et al*., 2012; Wetzel *et al*., 2016).

| Cytokine | Genotype | Cytokine secretion (pg) after |  |  |
| --- | --- | --- | --- | --- |
|  |  | High stimulation | Low stimulation | Con A stimulation |
| IL-1β | WT | 1,226 ± 91 | 864 ± 86 | 1,160 ± 132 |
| | *Syk*<sup>-/-</sup> | 604 ± 55 | 735 ± 51 | 750 ± 55 |
|  | *p* | 0.000039 | 0.19 | 0.0068 |
| IFN-γ | WT | 13,044 ± 3,307 | 9,950 ± 1,344 | 16,295 ± 3,659 |
| | *Syk*<sup>-/-</sup> | 16,006 ± 3,383 | 12,084 ± 2,146 | 14,059 ± 2,088 |
|  | *p* | 0.56 | 0.45 | 0.59 |
| IL-4 | WT | 1,094 ± 146 | 1,093 ± 213 | 1,168 ± 97 |
| | *Syk*<sup>-/-</sup> | 1,485 ± 240 | 1,237 ± 172 | 1,631 ± 364 |
|  | *p* | 0.20 | 0.60 | 0.30 |
| IL-10 | WT | 973 ± 292 | 1,666 ± 516 | 1,285 ± 433 |
| | *Syk*<sup>-/-</sup> | 1,547 ± 428 | 990 ± 241 | 2,091 ± 656 |
|  | *p* | 0.30 | 0.22 | 0.35 |
| IL-13 | WT | 824 ± 262 | 475 ± 104 | 396 ± 76 |
| | *Syk*<sup>-/-</sup> | 596 ± 119 | 498 ± 75 | 616 ± 84 |
|  | *p* | 0.41 | 0.85 | 0.075 |
| IL-17 | WT | < 156 | < 156 | < 156 |
|  | *Syk*<sup>-/-</sup> | < 156 | < 156 | < 156 |
| TNF-α | WT | < 124 | < 124 | < 124 |
|  | *Syk*<sup>-/-</sup> | < 124 | < 124 | < 124 |

Based on our data that SYK activity is required for *Leishmania* uptake by macrophages, we hypothesized that there would be fewer parasites contained within lesions of infected *Syk^flox/flox^* LysM Cre+ mice. Indeed, at experimental termination (wk 18), we found that lesions in *Syk^flox/flox^* LysM Cre+ mice contained fewer parasites than WT mice (Fig. 6B).

Next, we tested whether inhibition of SYK with entospletinib decreased lesion size and parasite burden. We found that entospletinib-treated mice developed smaller lesions than diluent/DMSO-treated mice over time (Fig. 6C). There were minimal effects on cytokine secretion when draining lymph nodes isolated from DMSO-treated infected mice were treated with entospletinib during cell culture (Table 2; Fig. S5F-I). The lone statistically significant result was an increase in IL-13 in some stimulated entospletinib-treated samples. Finally, we demonstrated that lesions in entospletinib-treated mice contained significantly fewer parasites than DMSO-treated mice at the termination of the experiment (wk 20, Fig. 6D).

**Table 2.** Culturing draining lymph nodes with entospletinib does not affect cytokine secretion. Isolated draining lymph nodes from 4 infected DMSO-treated mice were cultured as described as in Table 1, except that either DMSO or 2 μM entospletinib was added to each of these 4 isolated draining lymph nodes during culture for 72 h. Cytokine ELISAs were performed on harvested supernatants as in Table 1. Mean ± SE shown, with *p* values calculated by Students *t*-test. The lone statistically significant result (IL-13, high stimulation) is the opposite of the result that would be expected if immunological effects facilitated healing in entospletinib-treated mice (Jones *et al*., 1998).

| Cytokine | Treatment | Cytokine secretion (pg) |  |
| --- | --- | --- | --- |
|  |  | High stimulation | Low stimulation |
| IFN-γ | DMSO | 10,274 ± 509 | 13,455 ± 3,689 |
| | Entospletinib | 14,140 ± 4,374 | 19,226 ± 5,478 |
|  | *p* | n.s. | n.s. |
| IL-4 | DMSO | 1,522 ± 239 | 1,183 ± 511 |
| | Entospletinib | 1,047 ± 222 | 832 ± 187 |
|  | *p* | n.s. | n.s. |
| IL-10 | DMSO | 1,631 ± 442 | 2,013 ± 820 |
| | Entospletinib | 2,664 ± 424 | 2,451 ± 657 |
|  | *p* | n.s. | n.s. |
| IL-13 | DMSO | 342 ± 34.5 | 330 ± 36.6 |
| | Entospletinib | 491 ± 20.3 | 452 ± 53.2 |
|  | *p* | 0.021 | n.s. |

In summary, we have demonstrated that SFK and Abl family kinases activate SYK to facilitate phagocytosis, the uptake of *Leishmania* by Mϕs, and the disease manifestations seen in experimental cutaneous leishmaniasis. In combination with our previous results (Wetzel *et al*., 2012; Wetzel *et al*., 2016), these findings suggest that preventing parasite uptake by host cells using kinase inhibitors could be a valid strategy for designing novel antileishmanial therapies.

## DISCUSSION

We have shown previously that host SFK and Abl family kinases promote *Leishmania* uptake by Mϕs (Wetzel *et al*., 2012; Wetzel *et al*., 2016). Our results here demonstrate that SYK also is necessary for maximal *Leishmania* promastigote and amastigote uptake by Mϕs. Activation of SYK by SFK and Abl family kinases in M<s facilitates *Leishmania* internalization. Finally, mice lacking SYK in monocytes or treated with the SYK kinase inhibitor entospletinib exhibit significantly reduced lesion severity and parasite burden when infected with *Leishmania amazonensis*. Our data lead us to propose a relationship between FcR signaling, SFKs, Arg, and SYK during *Leishmania* amastigote uptake. Upon FcR ligation by amastigotes, Hck, Fgr, and Lyn are activated. These SFKs phosphorylate and activate Arg kinase, which phosphorylates and activates SYK, which leads to activation of CrkII, actin polymerization at the site of cell entry and formation/closure of the phagocytic cup (Fig. 6E).

Engagement of FcR stimulates downstream responses important for host immunity and receptor-mediated phagocytosis. Our results demonstrate that FcR-mediated signals permitting *Leishmania* amastigote uptake are relayed to SYK. SYK previously was shown to facilitate phagocytosis after opsonized particles bind FcR on phagocytic cells (Crowley *et al*., 1997). Reductions in IgG-mediated phagocytosis by *Syk^-/-^* BMDM were previously described to be secondary to defects in phagocytic cup formation and closure, rather than defects in Mϕ activation (Crowley *et al*., 1997), which is supported by our data. The effects of the novel SYK inhibitor, entospletinib, on phagocytosis had not been explored to date. We find that fewer IgG-coated beads or *Leishmania* amastigotes are internalized by entospletinib-treated M< or M< deficient in SYK. Neither condition changes the numbers of beads or parasites bound to Mϕ. The role of SYK in C3bi-mediated phagocytosis has been somewhat controversial (Kiefer *et al*., 1998; Tohyama and Yamamura, 2006; Walbaum *et al*., 2021). Our evidence supports SYK as a major contributor to efficient C3bi-mediated phagocytosis. Importantly, we also establish that SYK promotes the uptake of *Leishmania* promastigotes.

Multiple signaling pathways are regulated by SYK during phagocytosis, including linker for activation of T cells (LAT, (Balagopalan *et al*., 2015)), phospholipase Cγ (PLCγ, (Karimi *et al*., 1999)), and phosphatidylinositol 3-kinase (PI3-K, (Yamamori *et al*., 2000)). In addition, PI3-K signaling has been specifically implicated in *Leishmania* uptake by phagocytic cells (Cummings *et al*., 2012). In combination with previous studies, our experiments indicate that the specific signaling events necessary for FcR-mediated *Leishmania* amastigote internalization are initiated by the activation of SFK, which proceed to phosphorylate Arg kinase, which in turn phosphorylates SYK. Our data with entospletinib suggest that actin polymerization within phagocytic cups is increased, consistent with our prior findings using M< lacking Abl family kinases (Wetzel *et al*., 2012). The pSYK antibody used here specifically recognizes phosphorylation on Tyr525/526, which are SYK autophosphorylation sites that are necessary for maximal activity. Based on our data, we presume that Abl family kinases are modulating the phosphorylation of these two residues during amastigote uptake. However, our data do not rule out the possibility that other sites on SYK may also be phosphorylated. The link between Abl family kinases and SYK is likely to be direct, as previous studies have demonstrated that purified Arg kinase phosphorylates SYK *in vitro* (Bonnerot *et al*., 1998; Greuber and Pendergast, 2012; Parsa *et al*., 2008). Similarly, SFK have been shown to phosphorylate Arg kinase *in vitro* (Mader *et al*., 2011; Plattner *et al*., 2004; Tanis *et al*., 2003). We explored whether the extracellular domains of PSGL-1, which is known to activate SYK signaling via interactions with FcR (Zarbock *et al*., 2008), also participated in this signaling cascade, but were unable to find evidence that this alternate receptor was relevant during *Leishmania* uptake. However, the methods we used would not address the possibility that signaling occurs through the intracellular domains of PSGL-1; additional experiments will be conducted for this purpose. In addition, this study, in combination with our prior work (Wetzel *et al*., 2012; Wetzel *et al*., 2016) demonstrates that activation of SFK, Abl family kinases and SYK after FcR binding results in CrkII activation during the uptake of *Leishmania* parasites. A number of other downstream mediators of SYK signaling have been published (Mocsai *et al*., 2010). Our future studies will directly test whether these mediators, or other kinases previously thought to facilitate phagocytosis or internalization of other pathogens, also participate in the uptake of *Leishmania*.

Because *Leishmania* parasites that remain extracellular within a mammalian host are thought to either die or be killed by the host immune system, we tested whether SYK deficiency would restrict disease manifestations in a mouse model of cutaneous leishmaniasis. Our studies indicate that *Leishmania* survival and pathogenesis in mice depends on SFK (Wetzel *et al*., 2016), Abl/Arg (Wetzel *et al*., 2012; Wetzel *et al*., 2016), and now SYK. We believe that the smaller lesions seen in SYK-deficient mice primarily occur due to the deficits we identified in parasite internalization by phagocytes. Consistent with this hypothesis, there are fewer parasites within the lesions of SYK-deficient mice or mice treated with entospletinib. Since our hypothesis is difficult to test directly, our conclusions in part stem from eliminating other possibilities. For example, we found that entospletinib did not kill *Leishmania* parasites directly, and we saw no defects in parasite survival within entospletinib-treated M<s.

Furthermore, we assessed whether the reductions in lesion size observed in SYK-deficient mice could be due to variations in the host immune response to leishmaniasis compared to control mice. For example, SYK deficiency is known to affect B and T cell development in mice and immune cell adhesion (Mocsai *et al*., 2010), which might adversely affect the host’s ability to control infections. However, our results suggest that such defects play a more limited role in our experimental system. The overall innate immune response to leishmaniasis is complex and dependent upon the infecting parasite species and the mouse strain used. In general, Th1 responses lead to lesion healing and Th2 responses result in lesion worsening (Jones *et al*., 1998). Our profiling of cytokines secreted by isolated lymph nodes after infection revealed no significant differences in Th1 versus Th2 responses between SYK-deficient and control mice. Lymph nodes from *Syk^flox/flox^* LysM Cre+ mice did exhibit defects in IL1-β secretion, which is known to be partly dependent on SYK (Callaway *et al*., 2015). However, we would expect that a decrease in IL1-β would worsen lesions based on prior studies (Lima-Junior *et al*., 2013). Similarly, we did not see differences in secretion of cytokines from lymph nodes isolated from infected mice that were incubated in entospletinib, save for an increase in IL-13 in some stimulated entospletinib-treated samples, which also should not mediate lesion healing (Jones *et al*., 1998). Therefore, we saw no differences in the immune response to leishmaniasis from our cytokine profiling that would explain the decreased lesion size in mice deficient in SYK or treated with entospletinib. However, further studies are needed to fully delineate changes in the immunological response to SYK inhibition in experimental murine leishmaniasis.

## CONCLUSION

In summary, our studies demonstrate that SYK is required for maximal *Leishmania* uptake by Mϕs and disease in mice. Our future work will continue to trace the signaling pathways necessary for internalization of this parasite by Mϕ, with an eye towards using inhibitors of these pathways as treatment for leishmaniasis. A host-targeted therapeutic strategy is likely to be less susceptible to the development of resistance. While our studies have thus far focused on *Leishmania*, our findings likely are more broadly applicable, as other intracellular pathogens also utilize and activate SYK to facilitate receptor-mediated uptake. For example, overexpression of membrane-targeted SYK in cells treated with Abl kinase inhibitors partially rescues the impairment in phagocytosis of *Francisella tularensis* (Parsa *et al*., 2008). Thus, there is potential to harness host kinases such as SYK as targets for a broad-spectrum antimicrobial agent against intracellular pathogens.

## MATERIALS AND METHODS

### Mice

The UT Southwestern Institutional Animal Care and Use Committee approved all animal protocols. *Syk^flox/flox^ LysM Cre+* and *Syk^flox/flox^ LysM Cre-* littermates were generated by crossing *LysM Cre*+ mice to *Syk^flox/+^* mice (all on a C57BL/6J background, which along with WT C57BL/6J mice, had been purchased from Jackson Labs (Bar Harbor, ME)). *Hck^-/-^ Fgr^-/-^ Lyn^-/-^, Arg^-/-^*and *Abl^flox/flox^ LysM Cre+* mice were generated as described previously (Wetzel *et al*., 2012). For all experiments, WT littermates were paired with the kinase mutant under study to limit any effects from variation in genetic backgrounds.

### Mammalian cell culture

RAW 264.7 cells (obtained from ATCC, Manassas, VA) were incubated in Dulbecco’s modified Eagle’s medium (DMEM) (ThermoFisher, Waltham, MA) plus 10% heat-inactivated fetal bovine serum (FBS, Gemini). Bone marrow-derived primary Mϕs (BMDM) were harvested from tibias of *Syk^flox/flox^ LysM Cre+, Syk^flox/flox^ LysM Cre-, Hck^-/-^ Fgr^-/-^ Lyn^-/-^,* WT, *Arg^-/-^,* and/or *Abl^flox/flox^ LysM Cre+* mice and confirmed as described (Wetzel *et al*., 2012). All cells were screened for *Mycoplasma* contamination (Wetzel *et al*., 2016).

### Leishmania culture

*L. amazonensis* strain IFLA/BR/67/PH8 promastigotes (obtained from Norma W. Andrews, University of Maryland, College Park, MD) were grown in Schneider’s Drosophila medium (Sigma, St. Louis, MO) with 15% heat-inactivated FBS and 10 μg/ml gentamicin (Sigma) (Wetzel *et al*., 2012). 7 day old cultures were used for uptake experiments to maximize yield of infective metacyclic promastigotes. *L. amazonensis* strain IFLA/BR/67/PH8 amastigotes were grown axenically (outside of mammalian cells) in supplemented Schneider’s media at pH 5 as described (Hodgkinson *et al*., 1996). All parasites were serially passaged through mice so that virulence would be maintained.

### Transgenic *Leishmania* strains

For intracellular survival assays, *L. amazonensis* expressing a bright monomeric green fluorescent protein, mNeonGreen, was used. mNeonGreen was cloned into the pLEXSY.hyg2 (Jenabioscience, Jena, Germany) expression vector with a hygromycin resistance marker, as previously described (Ullah *et al*., 2020). pLEXSY.hyg2 containing mNeonGreen gene (pLEXSY.hyg2-mNeonGreen) was transfected into *L. amazonensis* promastigotes using the Human T-Cell Nucleofector kit and the Amaxa Nucleofector electroporator (program U-033, Lonza, Basel, Switzerland), for integration into the 18S rRNA locus within the nuclear DNA. Following transfections, promastigotes were allowed to grow for 24 h at 26 °C, then selected with 100 μg/mL hygromycin in Schneider’s Drosophila medium. Clones were isolated via limiting dilution, and the cultures were subsequently maintained in Schneider’s Drosophila medium supplemented with 100 μg/mL hygromycin. To maintain virulence, *L. amazonensis*-mNeonGreen expressing parasites were passaged regularly in C57BL/6 mice, as described previously (Wetzel *et al*., 2016).

### Phagocytosis assays

BMDM and RAW 264.7 cells were plated at > 50% confluence and starved overnight in macrophage-colony stimulating factor (M-CSF)-starved or serum-free media. For drug experiments, except where indicated, compounds were obtained from LC Laboratories (Woburn, MA). Mϕ were incubated in 3.3 μM imatinib, 1 μM SU6656, 1 μM entospletinib, or 0.1% DMSO (Sigma) for 2 h. For C3bi-coated bead internalization only, M< were also pre-activated with PMA (Sigma) for 1 h. As described (Wetzel *et al*., 2012; Wetzel *et al*., 2016), 2 μm latex green beads were incubated with human IgM (I-8260, Sigma), then in rabbit anti-human IgM (Sigma 270A, IgG-coated beads) or fresh mouse serum (C3bi-coated beads). 10 beads/ Mϕ were added to Mϕ for 30 m at 37 °C. To distinguish internal from external beads, samples were fixed in 3% formaldehyde for 15 m, blocked in 2% BSA (no permeabilization) (Wetzel *et al*., 2004; Wetzel *et al*., 2003; Wetzel *et al*., 2005), incubated in rabbit-anti human IgM and Hoescht 33258 (Sigma), and then incubated in Alexa-Fluor-564-conjugated goat anti-rabbit IgG. Samples were visualized using a Cytation 5 imager (BioTek, Santa Clara, CA) and analyzed by an observer blinded to condition. Representative images for beads were collected on the Cytation 5 at 40X and processed linearly in Adobe Photoshop (version 13.0.6) (Ullah *et al*., 2020). The phagocytic index (PI) was calculated as the number of beads internalized per 100 M<, and experimental samples were normalized to controls. The total number of beads per 100 M< was also calculated, with experimental samples were normalized to controls. At least 100 M< and 100 beads were counted from multiple imaged fields per condition. All experiments had at least 3 biological replicates performed, and means were calculated. Each biological replicate was then normalized so that the control was set to 100% for each experiment. The experimental category shown represents a percentage of this control value with standard errors. A one-sample *t-*test was used for statistical analysis of these results (Wetzel *et al*., 2012; Wetzel *et al*., 2016).

### *Leishmania* uptake assays

RAW 264.7 cells or BMDMs were treated with DMSO/PMA/inhibitors as above. To allow C3bi opsonization, we incubated metacyclic promastigotes in murine serum for 1 h. To allow IgG opsonization, we incubated amastigotes with anti-P8 primary IgG1 antibody (Pan and McMahon-Pratt, 1988) for 1 h. Opsonized parasites were incubated with M< at 10:1 promastigotes per Mϕ or 2:1 amastigotes per Mϕ (Wetzel *et al*., 2012; Wetzel *et al*., 2016). Samples were fixed with 3% formaldehyde, blocked in 2% BSA in PBS, incubated in mouse anti-gp46 antibody (promastigotes) or mouse anti-P8 antibody (both supplied by Diane McMahon-Pratt), then incubated in Alexa-Fluor-568 goat-anti-mouse secondary antibody (A10037, ThermoFisher) (Wetzel *et al*., 2012). After permeabilization, parasites were incubated in their original primary antibody, then Alexa-Fluor-488 donkey anti-mouse secondary antibody (A21202, ThermoFisher) and Hoescht 33258 (Sigma). Visualization, analysis, and statistics were performed as described above for bead phagocytic assays (Wetzel *et al*., 2012; Wetzel *et al*., 2016). Representative images for parasites were collected on a Zeiss LSM 880 inverted confocal Airyscan microscope at 40X (Ullah *et al*., 2020). Shown are maximal intensity projections constructed from full Z-thickness stacks through parasites/Mϕ using Image J (1.52a, http://imagej.nih.gov/ij) and processed linearly in Adobe Photoshop (version 13.0.6) (Ullah *et al*., 2020).

For PSGL-1 experiments, amastigotes were incubated with anti-P8 primary IgG1 antibody (Pan and McMahon-Pratt, 1988) for 1 h. RAW 264.7 cells were pre-treated with anti-mouse CD162 (PSGL-1) monoclonal antibody (4RA10; 12-1621-80, ThermoFisher), entospletinib, both, or equal amounts of vehicle (DMSO) for 1 hr. Antibody was employed at 1:2500, 1:1000, 1:500, 1:250, and 1:100, and entospletinib was applied at 1 uM. Opsonized parasites were incubated with RAW 264.7 cells at 10:1, respectively. Samples were fixed with 4% formaldehyde, blocked in 5% BSA/PBS, incubated in mouse anti-P8 antibody, then incubated in Alexa-Fluor-568 goat-anti-mouse secondary antibody (A10037, ThermoFisher) (Wetzel et al., 2012). After permeabilization with 0.1% Triton-X114, parasites were incubated in their original primary antibody, then Alexa-Fluor-488 donkey anti-mouse secondary antibody (A21202, ThermoFisher) and Hoescht 33258 (Sigma). Visualization was performed on an INCell Analyzer 6000™ (GE Healthcare, Inc.) microscope at 40x in the UTSW High Throughput Screening Core facility. For Fig. S3C, images were processed through a CellProfiler pipeline (https://cellprofiler.org). 3-6 technical replicates were performed for each condition and 3 separate biological replicates were combined to provide the data shown. Statistics were performed as above for phagocytic assays (Wetzel et al., 2012; Wetzel et al., 2016). Fig. S3D was analyzed as indicated under the “Phagocytosis assays” and “Leishmania uptake assays” headings.

### Viability assays

*L. amazonensis* promastigotes, axenic amastigotes, and RAW 264.7 cells were cultured as above, added to 96 well plates containing dilution series of relevant compounds, and incubated for 72 h before measuring viability using alamarBlue® (10%, ThermoFisher) (Ullah *et al*., 2020). Plates were read at 6 h with a BioTek Synergy H1 plate reader (530 nm excitation, 570 nm emission) (Ullah *et al*., 2020). To assess for parasite survival, 10 *L. amazonensis* promastigotes/Mϕ were added to G-CSF starved WT BMDMs and incubated for 4 h. After 5 washes with PBS, DMSO or 1 μm entospletinib was added in media with G-CSF. Samples were analyzed after 72 h by microscopy using the Cytation 5 imager, and total numbers of amastigotes/Mϕ were calculated by a blinded observer using the methods described above for phagocytosis assays.

### Immunoblotting

To measure phosphorylation of SYK or CrkII, RAW 264.7 cells were incubated overnight in serum-free media with experiments performed at ∼70% confluence. To initiate experiments, RAW cells were pre-incubated in 3.3 μM SU6656, 3.3 μM imatinib, 1 μM entospletinib, 50 nM DPH or DMSO for 2 h. IgG-opsonized amastigotes were added to adherent starved RAW 264.7 cells (at a ratio of 20:1) for 30 m (Pan and McMahon-Pratt, 1988). Cells were lysed with Pierce Radioimmunoprecipitation (RIPA) buffer (cat. no. 89901, Thermo Scientific, Rockford, IL) with protease and phosphatase inhibitors, and protein concentrations were assessed using bicinchoninic acid (BCA) (cat. no. 23225, Thermo Scientific). For representative images, equivalent protein amounts were loaded on 10% SDS-PAGE gels, transferred to nitrocellulose membranes, and probed with antibodies against phosphorylated Syk (pSyk) (Tyr525/526, cat. no. 2710S, Cell Signaling, Danvers, MA) at 1:500, or Syk (cat. no. 2712S, Cell Signaling) at 1:500. Immunoblotting with rabbit GAPDH (cat. no. 2118L, Cell Signaling) at 1:1000 was used as a loading control. For analysis, relative amounts of pSyk or pCrkII were compared with Image J analysis software and were normalized to Syk or CrkII (membranes were stripped of pSyk or pCrk and reprobed for Syk or Crk). Imaging was performed by using a phosphorimager (ImageQuant LAS 4000, GE, Boston, MA). Each experiment was performed 4 times. Two-way ANOVA was used to determine statistical significance.

### Murine infections

8 week old female C57BL/6 mice were infected in the dorsal side of the right hind foot with 1 X 10^6^ metacyclic promastigotes (isolated via Percoll gradient, Sigma) per mouse; 10 mice were used per experimental condition. All animals surviving until termination were included in analysis. For entospletinib-treated mice, 3 experiments were conducted. Mice were provided 100 mg/kg/d of entospletinib or the diluent (0.1 % DMSO) in their drinking water at pH 5.5, starting 4 days prior to the experiment and continuing through termination (Wetzel *et al*., 2016). For *Syk^flox/flox^ LysM Cre+* mice, 3 independent experiments were conducted. Lesion size was measured every other week with calipers by a condition-blinded investigator, and infected/uninfected foot thickness ratios were calculated and compared to time of infection by ANOVA. The number of parasites in lesions were determined at the time of experiment termination (between 18 and 20 wks post-infection) (Soong *et al*., 1997).

Lymph nodes from infected mice were harvested for cytokine profiling as described (Wetzel *et al*., 2012; Wetzel *et al*., 2016). In brief, cells from lymph nodes were plated and stimulated with varying concentrations of promastigote lysates (high stimulation = 2.5 X 10^6^ parasites; low = 5 X 10^5^ parasites) or concanamycin A (CON A) (5 μg/ml, Sigma); supernatants were harvested at 72 h. A negative (unstimulated control) was employed to ensure that cytokines released were secondary to stimulation, and levels of all measured cytokines were found to be negligible compared to background. Levels of IL-1β, IL-4, IFN-γ, IL-10, IL-13, and IL-17 were assessed by ELISA and compared to background (unstimulated) levels.

## Abbreviations

BMDM: bone marrow-derived macrophages
C3bi: terminal component of complement
CR3: complement receptor 3
FcR: Fc receptor
h: hour
IgG: immunoglobulin G
LPS: lipopolysaccharide
Mϕ: macrophage
m: minutes
SE: standard error
p-: phosphorylated
SFK: Src family kinases
SYK: spleen tyrosine kinase

## Acknowledgements

We thank the parasitology group at UT Southwestern (Drs. James J. Collins, Margaret A. Phillips, and Michael L. Reese), as well as members of the UT Southwestern Departments of Biochemistry, Microbiology, Pharmacology, and Pediatrics for valuable discussions and use of common equipment. Gina M. Aloisio, Lauren T. Callaghan, Ivan Luu, Hanspeter Niederstrasser, T. Michael Oladugba, and Catherine L. Trice provided expert technical assistance. Dr. Naomi S. Morrissette (UC Irvine) supplied helpful comments on the manuscript, and Drs. Anthony J. Koleske and Diane McMahon-Pratt (both from Yale University) offered valuable advice as this project was conceived. Antileishmanial antibodies were donated by Dr. McMahon-Pratt. *Arg^-/-^, Abl^flox/flox^*, and *Arg^-/-^ Abl^flox/flox^* mice were provided by Dr. Koleske. *Hck^-/-^ Fgr^-/-^ Lyn^-/-^* mice were provided by Shaoguang Li. Confocal microscopy was performed in the UT Southwestern Live Cell Imaging Facility.

## Author contributions

I.U., U.B., A.D., and D.M.W. designed and performed experiments, analyzed data, and wrote the manuscript. R.M.K., F.T.H.K., E.T.M., L.M.B, E.L.R., and J.M.B. performed experiments and analyzed data.

## Data availability

This manuscript was uploaded to Biorxiv as a preprint prior to publication (BIORXIV/2022/513575). All data referenced in this manuscript is available upon publication within the main body or supplementary data sections or on Biorxiv.

---

**Fig. S1.**
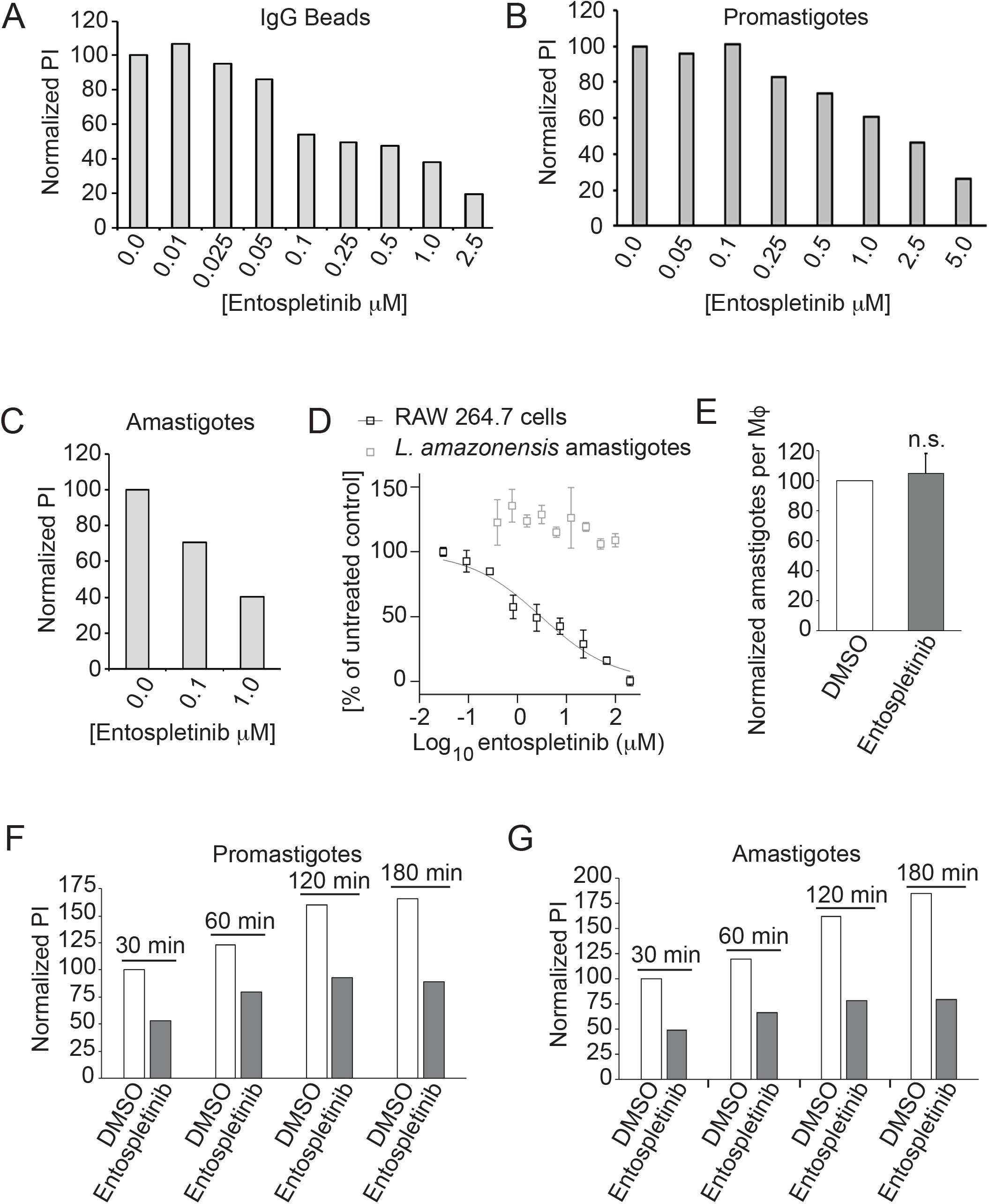
Titrations of entospletinib dosage and effects over time. (A) Titration of entospletinib’s effects on IgG-coated bead uptake by RAW 264.7 cells. Mϕs were treated with DMSO (shown as 0.0 μM entospletinib) or increasing concentrations of entospletinib for 2 h and incubated with IgG-coated beads for 30 m. Shown is one representative experiment of 2 experiments. Data normalized to DMSO category (PI = 100%). (B) Titration of entospletinib’s effects on promastigote uptake by RAW 264.7 cells. Mϕs were treated with increasing concentrations of entospletinib or DMSO for 2 h and incubated with IgG-coated amastigotes for 30 m. Shown is one representative experiment of 2 experiments. (C) Titration of entospletinib’s effects on amastigote uptake by RAW 264.7 cells. Mϕs were treated with DMSO, 0.1 μM, or 1 μM entospletinib and incubated with IgG-coated amastigotes for 30 m. Shown is one representative experiment of 2 experiments. (D) Entospletinib log concentration-response curves for RAW 264.7 cells (black) and axenic *L. amazonensis* amastigotes (grey) for one representative experiment of 3 biological replicate experiments. Triplicate technical replicates were used. Cells were incubated for 72 h in the concentrations of entospletinib shown. (E) Treatment with 0.5 μM entospletinib has no effects on *L. amazonensis* amastigote survival inside RAW 264.7 cells. Mϕs were allowed to internalize *L. amazonensis* amastigotes expressing mNeonGreen, and then, after 24 h, they were incubated with DMSO or 0.5 μM entospletinib for 72 h. Shown is the mean number of *L. amazonensis* amastigotes per 100 entospletinib-treated Mϕ, normalized to DMSO-treated Mϕ (100%), ± SE. n = 3 separate biological replicates. (F) The relative decrease in promastigote PI in entospletinib-treated RAW 264.7 cells does not change after long drug incubations. Mϕs were treated with 1 μM entospletinib or DMSO for 2 h and incubated with C3bi-coated promastigotes for 30, 60, 120, or 180 m. Data normalized to DMSO, 30 m category. Shown is one representative experiment of 2 experiments. (G) The relative decrease in amastigote PI in entospletinib-treated RAW 264.7 cells does not change after long drug incubations. Experiment performed as in (F), except that M<s were incubated with IgG-opsonized amastigotes. Shown is one representative experiment of 2 experiments.

**Fig. S2.**
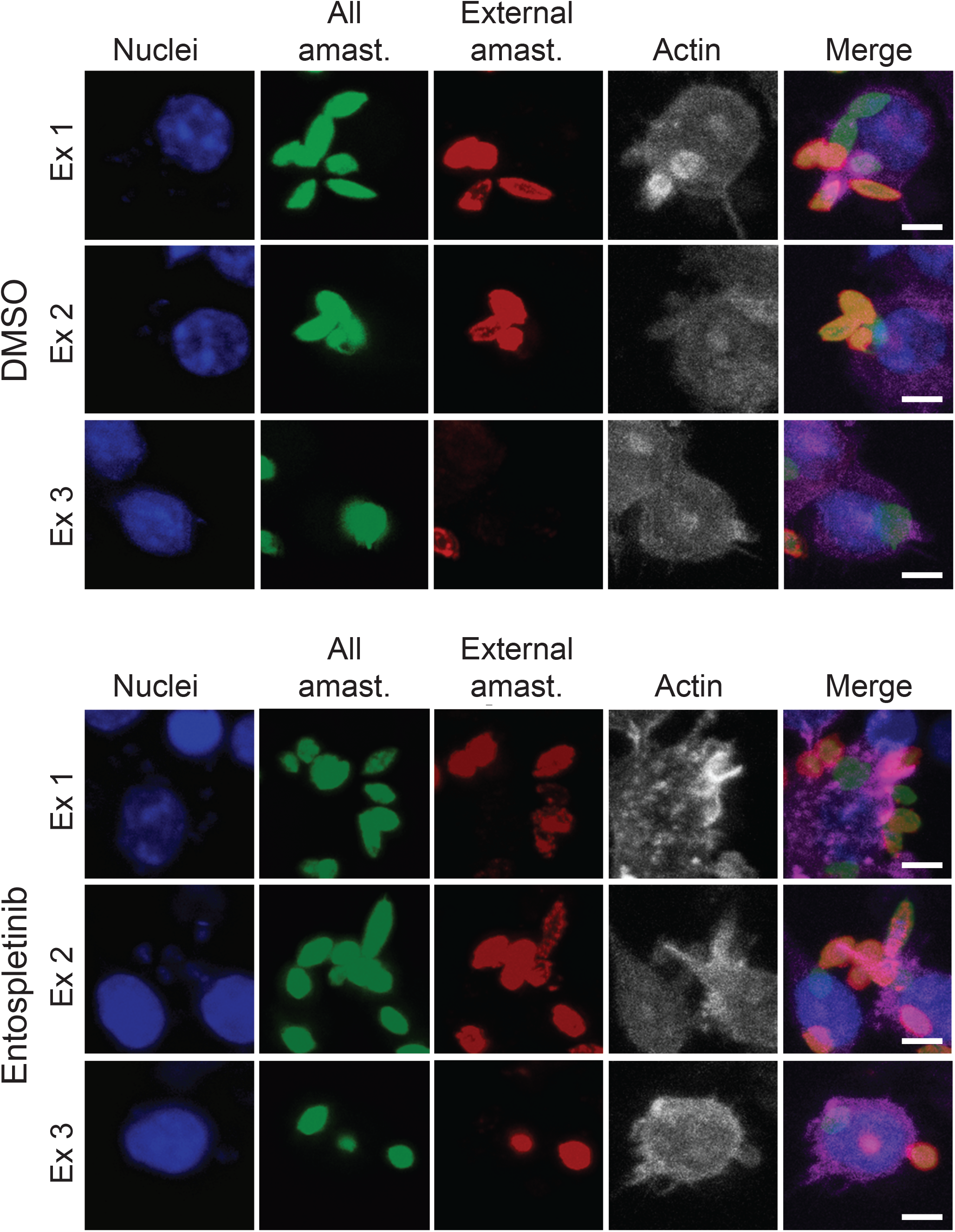
Representative phagocytic cups in DMSO vs entospletinib-treated M<s taking up *Leishmania* amastigotes. Shown are three separate examples of phagocytic cups in DMSO versus entospletinib-treated RAW 264.7 cells that are in the process of internalizing amastigotes. From left to right: M< nuclei (blue, Hoescht); all amastigotes (green; p8); external amastigotes (red); actin (white/pink; phalloidin). Scale bar = 2 μm. Circular phagocytic cups like those seen in example 1 in the DMSO-treated M<s were not seen in entospletinib-treated M<s, consistent with prior literature suggesting that there are defects in cup closure in *SYK^-/-^* M<s (Crowley *et al*., 1997). Actin staining also appears brighter in entospletinib-treated M<s than DMSO-treated M<s (quantified in Fig. S3B).

**Fig. S3.**
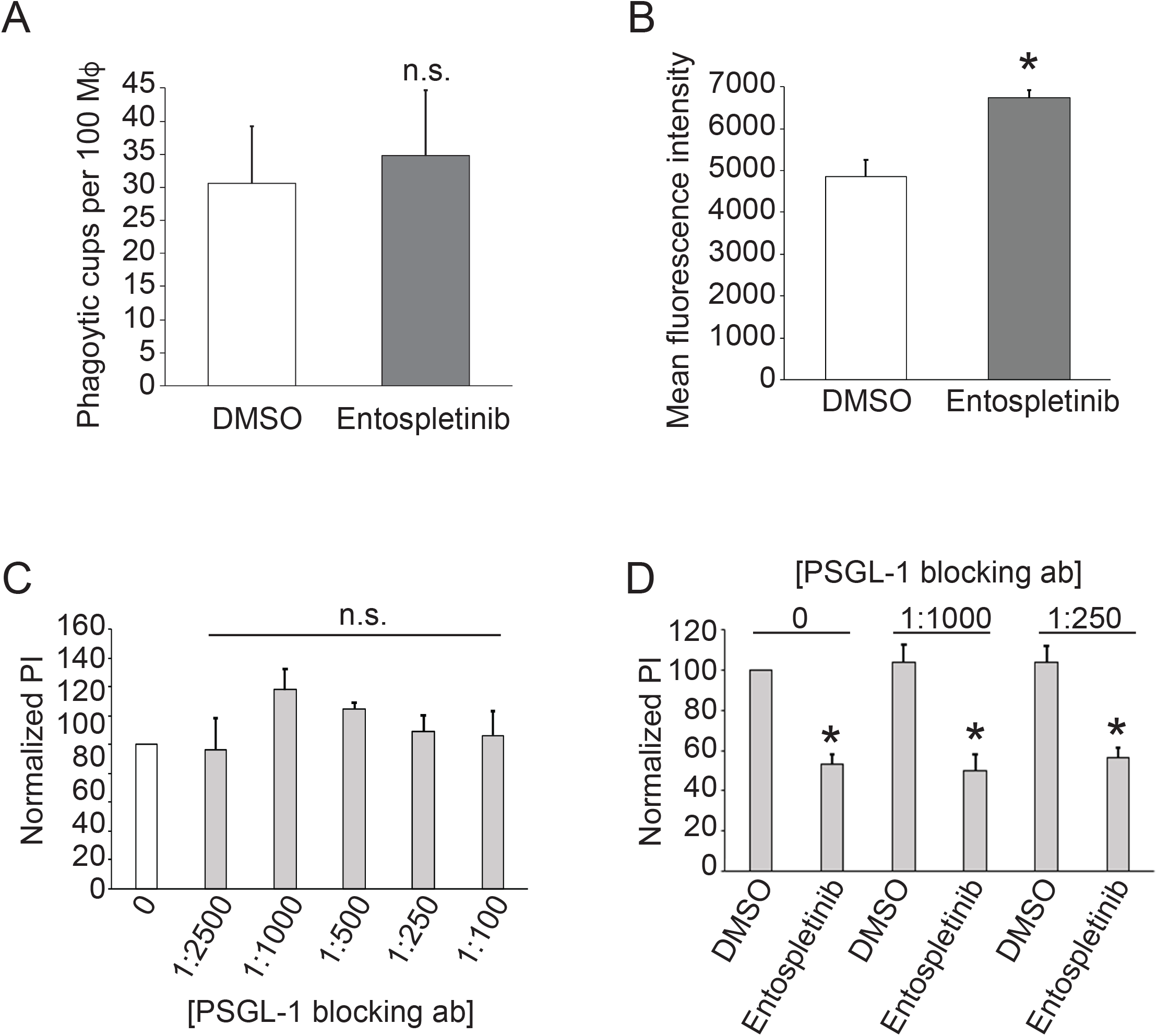
Mechanisms of SYK signaling during *Leishmania* uptake. (A) Despite differences in PI, the number of phagocytic cups seen between entospletinib-treated and DMSO-treated RAW 264.7 cells incubated with amastigotes is the same. This result is consistent with prior literature documenting defects in phagocytic cup closure (not formation) in *Syk^-/-^*phagocytes. (B) Actin in phagocytic cups shown in Fig. S2 is brighter in entospletinib-treated Mϕs than in DMSO-treated Mϕs. Relative fluorescence intensity was quantified in ImageJ by an observer blinded to experimental category. Shown is the mean intensity per field among at least 5 fields per category ± SE. *, p< 0.05 by *t*-test. (C) Incubation with 4RA10, an antibody to P-selectin glycoprotein ligand 1 (PSGL-1), does not affect uptake of *Leishmania* amastigotes by Mϕs. Mϕs were treated with increasing concentrations of 4RA10 for 2 h and incubated with IgG-coated amastigotes for 30 m to allow internalization as described above. Shown is the mean PI ± SE from 3 biological replicates. All categories non-significant by ANOVA. (D) The entospletinib-induced reduction in amastigote PI is not affected by 4RA10. Mϕs were treated with increasing concentrations of 4RA10 and DMSO vs 1 μM entospletinib for 2 h, then incubated with IgG-coated amastigotes for 30 m to allow internalization as described above. Shown is the mean PI ± SE from 3 biological replicates. *, p< 0.05 by ANOVA.

**Fig. S4.**
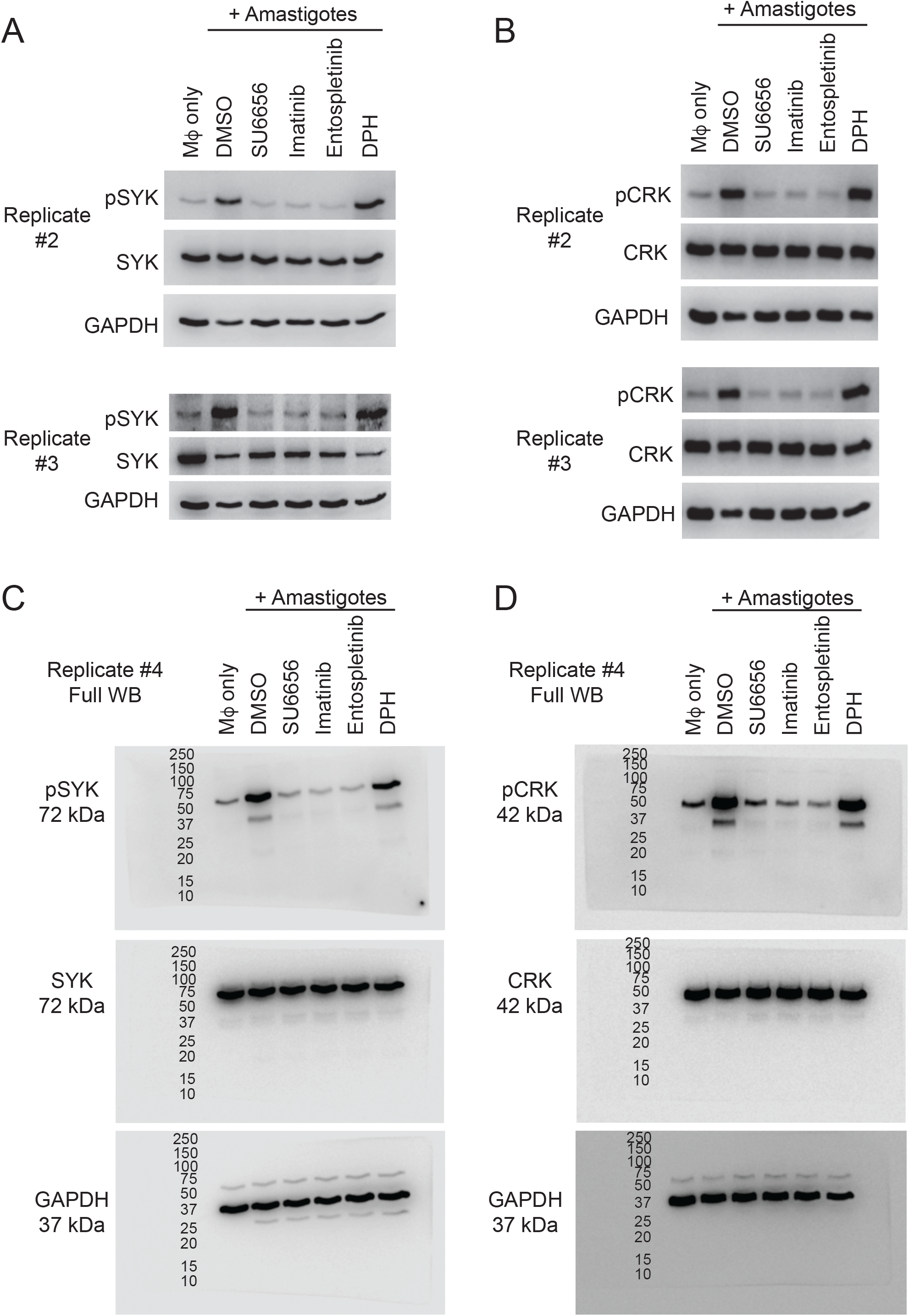
Western Blots. All Western blots quantified in Figure 5 are shown. (A) Biological replicates 2 and 3 of Western blots quantified for pSYK calculations. (B) Biological replicates 2 and 3 of Western blots quantified for pCRK calculations. (C) Full Western blot for biological replicate 4 used for pSYK calculations. (D) Full Western blot for biological replicate 4 used for pCRK calculations.

**Fig. S5.**
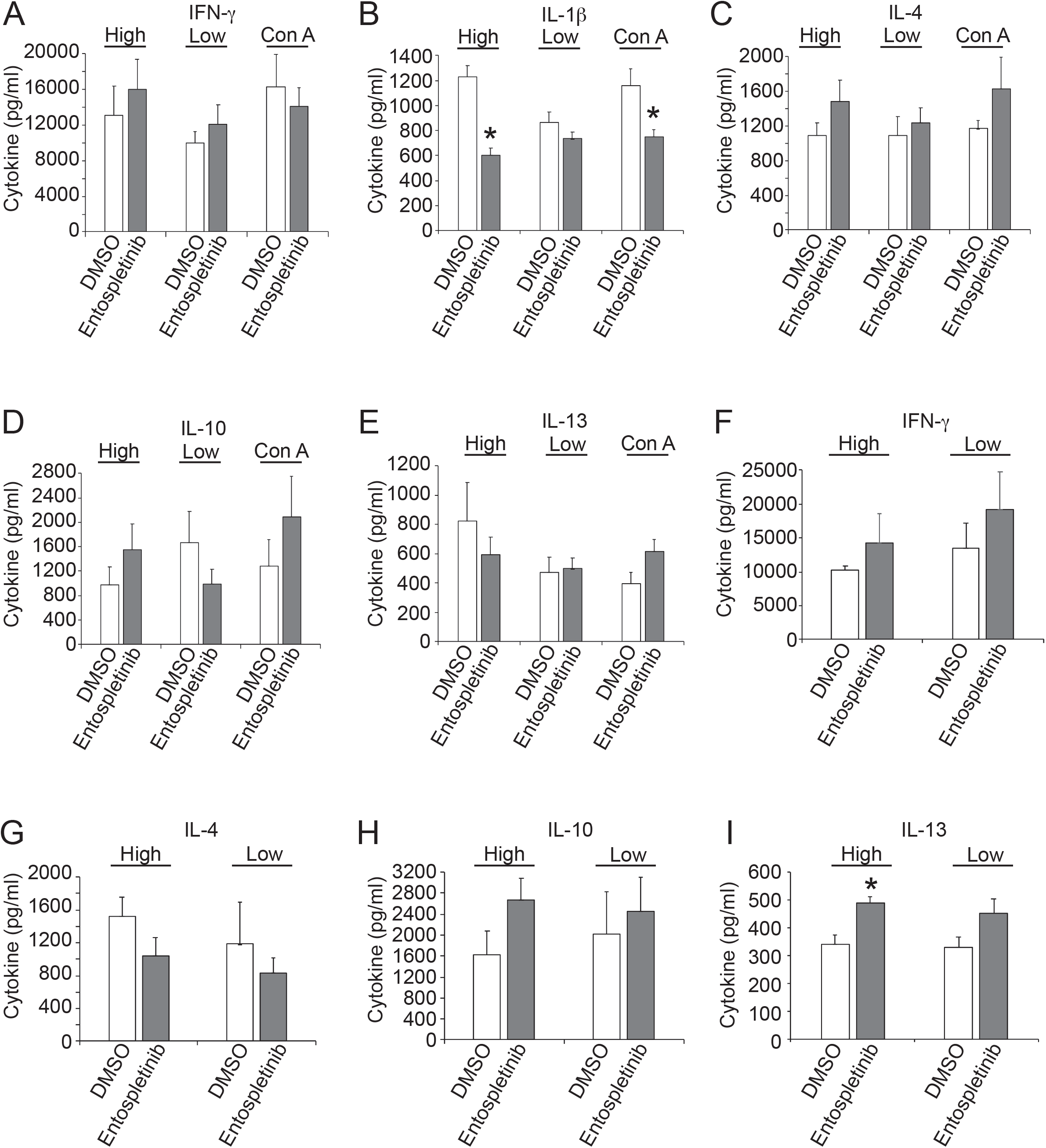
Graphical representations of cytokine secretion data. (A-E) Graphs of data shown in Table 1, separated out by cytokine. A = IFN-γ, B= IL-1ϕ, C = IL-4, D = IL-10, E = IL-13. (F-I) Graphs of data shown in Table 2, separated out by cytokine. F = IFN-γ, G = IL-4, H = IL-10, I = IL-13. *, p< 0.05 by *t*-test.

**Table S1.**
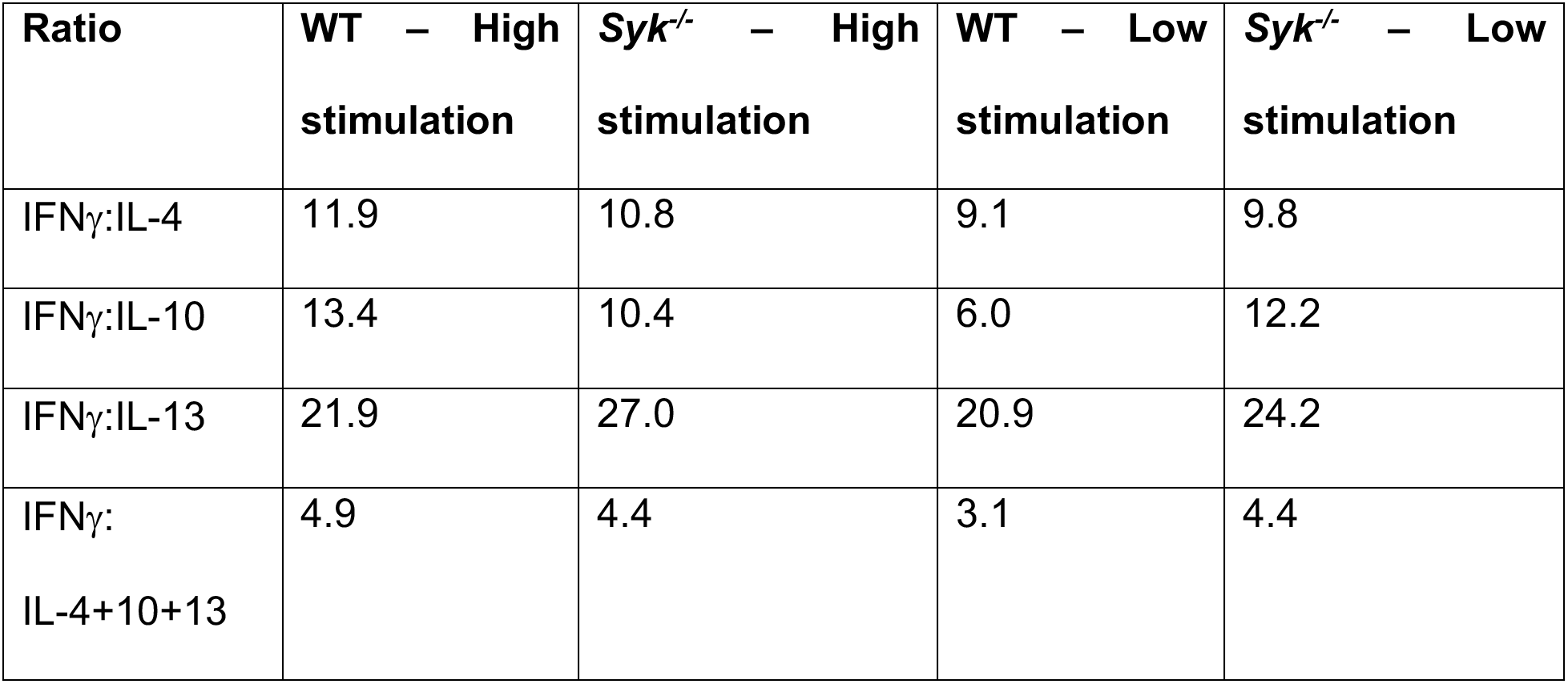
Cytokine secretion is not skewed towards a Th1 or Th2 response during *L. amazonensis* infection of *Syk^flox/flox^* LysM Cre+ mice compared to WT mice. Ratios of IFNγ:IL-4, IFNγ:IL-10, IFNγ:IL-13, and IFNγ:IL-4+10+13 were calculated from the data shown in Table 1.

| Ratio | WT – High stimulation | *Syk<sup>-/-</sup>* – High stimulation | WT – Low stimulation | *Syk<sup>-/-</sup>* – Low stimulation |
| --- | --- | --- | --- | --- |
| IFNγ:IL-4 | 11.9 | 10.8 | 9.1 | 9.8 |
| IFNγ:IL-10 | 13.4 | 10.4 | 6.0 | 12.2 |
| IFNγ:IL-13 | 21.9 | 27.0 | 20.9 | 24.2 |
| IFNγ:<br>IL-4+10+13 | 4.9 | 4.4 | 3.1 | 4.4 |

